# Leveraging functional genomic annotations and genome coverage to improve polygenic prediction of complex traits within and between ancestries

**DOI:** 10.1101/2022.10.12.510418

**Authors:** Zhili Zheng, Shouye Liu, Julia Sidorenko, Loic Yengo, Patrick Turley, Alireza Ani, Rujia Wang, Ilja M. Nolte, Harold Snieder, Lifelines Cohort Study, Jian Yang, Naomi R Wray, Michael E Goddard, Peter M Visscher, Jian Zeng

**Affiliations:** Institute for Molecular Bioscience, The University of Queensland, Brisbane, Queensland, Australia; Program in Medical and Population Genetics, Broad Institute of Harvard and MIT, Cambridge, Massachusetts, USA; Stanley Center for Psychiatric Research, Broad Institute of Harvard and MIT, Cambridge, Massachusetts, USA; Analytic and Translational Genetics Unit, Massachusetts General Hospital, Boston, Massachusetts, USA; Center for Economic and Social Research, University of Southern California, Los Angeles, CA, USA; Department of Economics, University of Southern California, Los Angeles, CA, USA; Department of Epidemiology, University of Groningen, University Medical Center Groningen, Groningen, Netherlands; Department of Bioinformatics, Isfahan University of Medical Sciences, Isfahan, Iran; School of Life Sciences, Westlake University, Hangzhou, Zhejiang, China; Westlake Laboratory of Life Sciences and Biomedicine, Hangzhou, Zhejiang, China; Queensland Brain Institute, The University of Queensland, Brisbane, Queensland, Australia; Faculty of Veterinary and Agricultural Science, University of Melbourne, Parkville, Victoria, Australia; Biosciences Research Division, Department of Economic Development, Jobs, Transport and Resources, Bundoora, Victoria, Australia

## Abstract

We develop a new method, SBayesRC, that integrates GWAS summary statistics with functional genomic annotations to improve polygenic prediction of complex traits. Our method is scalable to whole-genome variant analysis and refines signals from functional annotations by allowing them to affect both causal variant probability and causal effect distribution. We analyse 28 traits in the UK Biobank using ∼7 million common SNPs and 96 annotations. SBayesRC improves prediction accuracy by 14% in European ancestry and by up to 33% in trans-ancestry prediction, compared to the baseline method SBayesR which does not use annotations, and outperforms state-of-the-art methods LDpred-funct, PolyPred-S and PRS-CSx by 12-15%. Investigation of factors affecting prediction accuracy identified a significant interaction between SNP density and annotation information, encouraging future use of whole-genome sequence variants for prediction. Functional partitioning analysis highlights a major contribution of evolutionary constrained regions to prediction accuracy and the largest per-SNP contribution from non-synonymous SNPs.

## Introduction

Polygenic scores (PGS) for complex traits (also known as polygenic risk scores (PRS) for common diseases) are playing increasingly important roles in research and medical applications of the fast-growing genomic data from genome-wide association studies (GWAS)^1^. PGS are used to provide evidence of polygenic adaptation of populations to different environments^2^, to explore putative causal relationships between traits^3^, to improve cost and efficiency of clinical trials^4^, and perhaps most importantly, to identify individuals with high genetic risk of complex diseases^5-10^, which provides opportunities for preventative medicine, early intervention, improved prognosis and personalised treatment^11-13^. However, the clinical application of PGS is currently limited by the modest prediction accuracy for most complex diseases. Furthermore, a substantial loss of prediction accuracy is observed when applying PGS across ancestries^14-20^.

Current PGS methods (see ref^21^ for a review) rely on estimating the effects of common SNPs that tag the causal variants by linkage disequilibrium (LD). The prediction accuracy then depends on the estimation accuracy of the SNP effects and the extent to which the LD in the GWAS population is the same as that in the target population. First, complex traits are affected by many common causal variants each with a vanishingly small effect size, likely due to the action of negative selection^22-24^. Therefore a very large sample size is required to accurately estimate their effects. Given the limited sample sizes for most diseases, it would be helpful to incorporate other information that are independent of the GWAS data. Second, although the parsimonious assumption that common causal variants are shared across ancestry groups has empirical support^20,25^, the LD between the causal variants and the SNP markers are likely to vary between populations^20^. Thus, to maximise prediction accuracy of PGS across people of different ancestries requires selection of the causal variants into the prediction equation rather than non-causal variants that tag the causal variants by LD.

Functional genomic annotations, e.g., the chromatin state and biological function of a genomic region, are informative regarding the locations and effect sizes of the causal variants underlying complex traits^26,27^. This information can be used to separate the likely causal SNPs from non-causal SNPs in high LD with them^28^ and hence improve polygenic prediction^15,29-32^. Harnessing functional annotations to improve prediction was first proposed in livestock genetics in a method called BayesRC^33^, a Bayesian method that incorporates biological priors for SNP effects but requires individual-level genotype and phenotype data and only allows discrete annotation categories. Recent methodological development in human genetics have allowed the integration of GWAS summary-level data with overlapping annotations for polygenic prediction, such as AnnoPred^30^, LDpred-funct^31^, MegaPRS^32^ and PolyPred^15^. However, there are limitations in these methods. First, most of these methods assume sparse genetic architecture by a mixture prior, which is superior to an infinitesimal assumption and is likely a better representation for many traits^23,34-36^, but consider only a subset of common variants, e.g., SNPs from a genotyping array or the HapMap3 panel^37^, due to computational feasibility. We caution that there is potentially a problem in modelling the annotation information when considering only a subset of SNPs, because SNPs that tag the causal variants by LD do not necessarily tag them by annotation, resulting in a mismatch between effect size and biological function of the variant (as illustrated in **Fig. 1a**). Second, these methods rely on the estimated per-SNP heritability enrichment for each annotation^38^ as input data, which can, supported by empirical results^33,39^, arise from either differences in the proportion of causal variants or in the distribution of effect sizes between annotation levels or categories (**Fig. 1b**), but none of the methods explicitly accounts for these two sources of information in the model. Third, all of them are stepwise weighting methods that require a tuning sample of individual-level data from the target population, which is not available for many traits.

**Figure 1.**
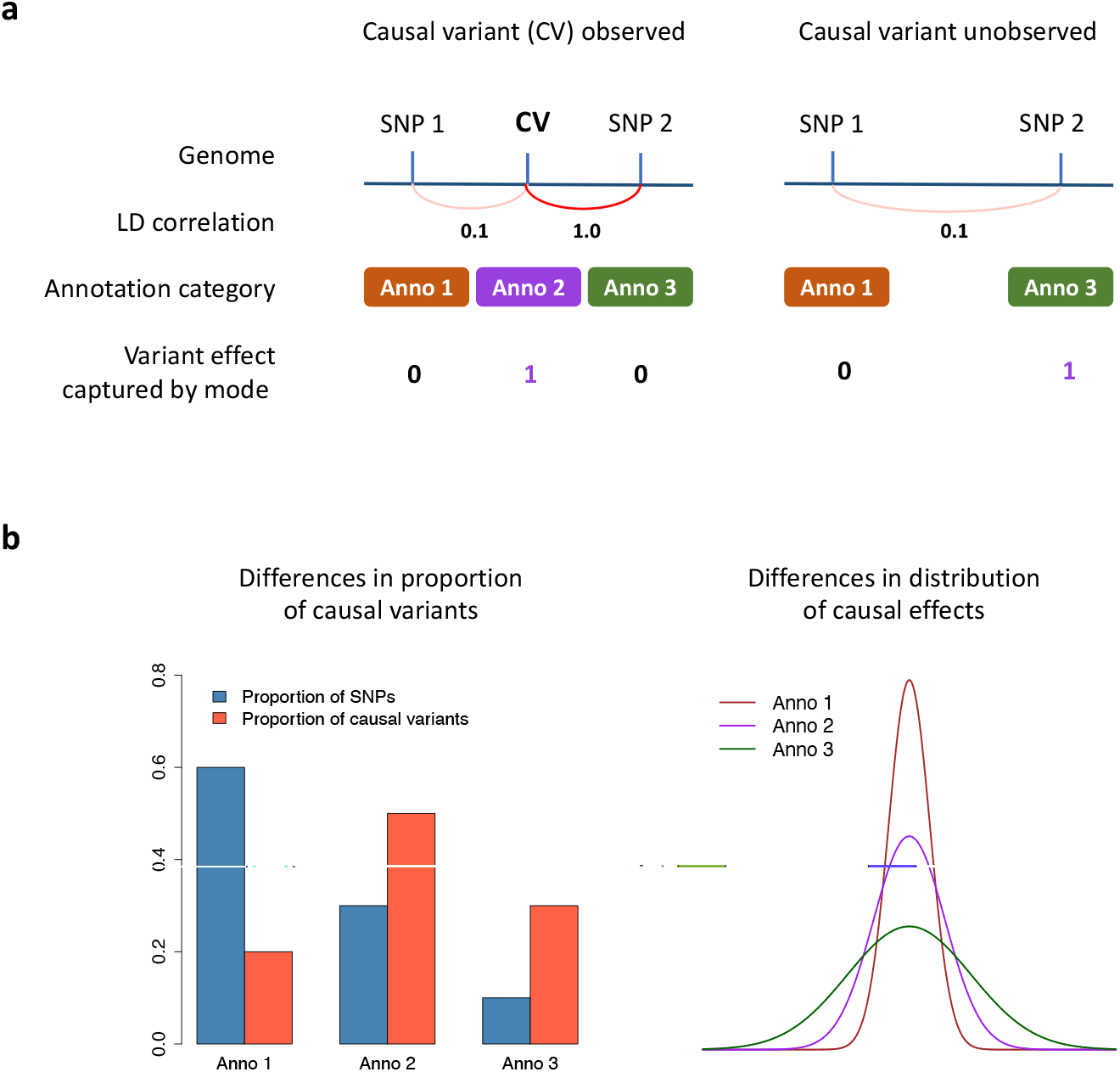
Characteristics of functional annotation data. a) Functional annotations provide orthogonal information that help to distinguish the causal variant from the SNP in perfect LD with it. However, when the causal variant is not observed, the SNP that tags the causal variants by LD captures the causal effect but does not share the same annotation with the causal variant, resulting in a mismatch between effect size and annotation category. b) Functional categories can differ in both the proportion of causal variants and the distribution of causal effect sizes, either of which can lead to an enrichment or depletion in per-SNP heritability in a functional category.

Here, we propose a new method, SBayesRC, that addresses these limitations by 1) analysing all imputed common SNPs simultaneously using an efficient algorithm, 2) refining the annotation information using a hierarchical multi-component mixture prior that allows annotation data to affect the probability of a SNP being causal as well as that of having a certain magnitude of causal effect size, and 3) using a full Bayesian learning machinery that estimates all parameters jointly from the data without a need of tuning. We apply our method to 28 complex traits including diseases using up to 10 million imputed common SNPs with 96 functional annotations (including discrete and continuous annotations), consider both within European ancestry and trans-ancestry prediction using UK Biobank^40^ and Lifeline^41^ datasets, and make comparisons with the best methods in the literature (**Table 1**). We also investigate factors that affect prediction accuracy when exploiting annotation data, and consider connections between the genetic architecture of functional categories and their contributions to prediction accuracy.

**Table 1.** Summary of prediction methods used in this study.

| Method | Features | Use annotations | Use all imputed SNPs | Require tuning data |
| --- | --- | --- | --- | --- |
| C+PT<sup>44</sup> | Non-parametric method based on GWAS marginal effect estimates of a subset of independent SNPs obtained by LD clumping and p-value thresholding. | No | No | Yes |
| LDpred2<sup>43</sup> | Bayesian mixture model that fits SNPs jointly with a point-normal prior. It is more powerful than C+PT by accounting for sparse genetic architecture and LD between SNPs. | No | No | Yes |
| LDpred-funct<sup>31</sup> | Stepwise method that starts with an infinitesimal model with functionally informed priors and then considers sparse genetic architecture empirically by regularizing SNP effects in bins of different magnitude using cross-validation. Compared to LDpred2, it incorporates functional annotations and analyses all imputed SNPs. | Yes | Yes | Yes |
| SBayesR<sup>35</sup> | Multi-component mixture model that assumes a finite mixture of normal distributions for SNP effects. It accounts for complex genetic architecture. All parameters in the mixture distribution are estimated from data. | No | No | No |
| SBayesRC (this work) | Extension of SBayesR to incorporate functional annotations and allow for fitting all imputed SNPs jointly. It considers differences between annotations in both the proportion of causal variants and the magnitude of causal effect sizes. | Yes | Yes | No |
| PolyPred-S<sup>15</sup> | Stepwise method that combines SNP effect estimates from a functionally informed fine-mapping approach<sup>50</sup> for 18M SNPs with those from SBayesR for 1M HapMap3 SNPs. It is flexible to the choice of method that generates the joint-effect estimates of HapMap3 SNPs, for which SBayesR is recommended when LD reference sample closely matches the GWAS sample. It is proposed mainly for cross-population prediction, and uses a sample from the target population for weighting the SNP effects. | Yes | Yes | Yes |
| PRS-CSx<sup>14</sup> | Multi-discovery method designed to improve cross-population prediction by jointly modelling summary statistics data from multiple ancestrally diverse populations. It borrows information across populations by a shared continuous shrinkage prior but still requires a tuning data to maximize the prediction accuracy in the target population. It does not consider functional annotations and is computationally challenging to fit more than 1M HapMap3 SNPs. | No | No | Yes |
| SBayesRC-multi (this work) | Extension of SBayesRC to combine the SNP effect estimates from multiple populations after running SBayesRC in each population separately. Similar to PRS-CSx, a tuning sample of individual-level data is used to derive the final predictors for the target population. However, unlike PRS-CSx, the GWAS summary statistics from different populations are not jointly modelled. | Yes | Yes | Yes |

## Results

### Method overview

SBayesRC extends SBayesR^35^ to incorporate functional annotations and allows joint analysis of all common SNPs in the genome. Similar to SBayesR, SBayesRC only requires summary statistics from GWAS (i.e., marginal SNP effect estimates, standard errors, and GWAS sample size) and LD correlations from a reference sample as input data. In addition to joint SNP effect estimates for deriving PGS, SBayesRC also generates the fine mapping Bayesian statistics of posterior inclusion probabilities (PIP) for SNPs as measures of trait associations, and estimates of functional genetic architecture parameters such as SNP-based heritability and polygenicity associated with the functional annotations.

The capability of simultaneously analysing all common variants is achieved by deriving a low-rank model based on the eigen-decomposition on quasi-independent LD blocks in the human genome^42^. In each LD block *k*, we transform the summary-data-based model (as used in SBayesR) into a new model where the joint effects of *m*_*k*_ SNPs are fitted to *q*_*k*_ linear combinations of marginal SNP effects (instead of *m*_*k*_ marginal SNP effects), with *q*_*k*_ being the number of top principal components that collectively explain more than a given proportion (*ρ*) of variance in the LD matrix (**Methods**; **Supplementary Fig. 1a**). In such a model, the dimension of the system of equations is *q*_*k*_ × *m*_*k*_, usually *q*_*k*_ is much smaller than *m*_*k*_, leading to considerably less memory consumption and faster computation (**Table 2**). For instance, when *ρ* = 99.5%, *q*_*k*_/*m*_*k*_ ≈ 0.2 on average across LD blocks for the 7.4 million common SNPs used in this study (**Supplementary Fig. 2**). Essentially, the low-rank model refines the signals in GWAS summary statistics by collapsing information from SNPs in high LD. This also leads to independent residuals for the transformed observables, which warrants a more robust algorithm by estimating the residual variance from the data (**Methods**).

**Table 2.** Computation resource required for different methods. Results are average values across traits using 4 CPU cores when multi-thread is supported for the method.

| Method (No. SNPs) | Runtime (hours) | Memory (GB) | Required Storage (GB) |
| --- | --- | --- | --- |
| SBayesRC (7M) | 9.5 | 75.1 | 130 |
| LDpred-funct (7M) | 6.0 | 120.6 | 40-50 per trait |
| PolyPred-S (7M) | 19.8<sup>1</sup> | 71.7 | 2,800 |
| LDpred2 (1M) | 5.5 | 53.4 | 43 |
| SBayesRC (1M) | 1.2 | 7.8 | 5.6 |
| SBayesR (1M) | 0.5 | 27.0 | 22 |
| PRS-CSx (1M) | 14.2<sup>2</sup> | 4.7 | 5.6 |

<sup>1</sup> In PolyPred-S, fine-mapping is the most time-consuming step and is suggested to run by blocks in parallel.
Here we used a single core and divided the runtime by 4 for comparison to others.
<sup>2</sup> Per data set runtime: total runtime / number of training data sets (= 2).

We use an annotation-dependent prior to better model the distribution of SNP effects, and learn both annotation parameters and SNP effects from the data. In SBayesR, SNP effects are assumed to follow a mixture of normal distributions with different variances, including a point mass at zero. This model accounts for a spectrum of genetic architecture from major-gene to infinitesimal architecture and allows the data to inform which is the best-fit for the trait of interest. It is, however, independent of genomic features, such as LD and minor allele frequency (MAF) patterns and biological functions of different genomic regions. In SBayesRC, the probability parameters of the mixture distribution are estimated from a linear model in which the individual SNP annotations are the independent variables. This describes the probability that SNPs are causal variants and the probability distribution of their effect sizes (**Methods**; **Supplementary Fig. 1b**). In other word, we allow the distribution of SNP effects to be dependent on annotations, and this model will better capture the causal effects if the distributions of effect sizes are truly different between annotations. Owing to the use of a generalised linear model for the annotation effects, both binary and quantitative annotations are accommodated irrespective of overlapping between them. Our method has been implemented in a user-friendly software *GCTB*^23^ and in a R package (**Code availability**).

### Genome-wide simulation based on real genotypes and annotation data

We first used simulation to calibrate our low-rank model which requires a specification of two parameters: the minimum width (cM) of a LD block after merging contiguous small quasi-independent LD blocks^42^, and the minimum proportion (*ρ*) of variance in the LD matrix explained by the selected principal components in each LD block. We simulated GWAS data using 1,154,522 SNPs on 328,501 individuals of European ancestry in the UKB and calculated the prediction accuracy in a hold-out sample of 14,000 individuals. We found that in general the prediction accuracy increased with the increasing minimum value of LD block width, reaching to a plateau at the width of 4cM (**Supplementary Fig. 3**). When *ρ* increased from 85% to 99.95%, prediction accuracy improved due to increased amount of LD information, which allowed SNPs to better track the causal variants, reaching a plateau at *ρ* = 99.5%. Thus, for the best computational performance and a stable prediction accuracy, we decided to use a minimum block width of 4cM and *ρ* = 99.5% in the subsequent analyses unless otherwise noted.

We then checked the robustness of our method to two common challenges in the real data analysis: 1) differences in LD between GWAS and LD reference datasets, and 2) unequal GWAS sample sizes across SNPs (**Supplementary Note**), in comparison to two state-of-theart methods using summary statistics, LDpred2^43^ and SBayesR^35^. For all methods, the prediction accuracies decreased when the LD reference sample size was too small relative to the GWAS sample size (large variation in LD by chance), but SBayesRC (no annotation used) preserved more prediction accuracy than the other methods (**Fig. 2a** and **Supplementary Fig. 4**). In an extreme case where African ancestry individuals were used to estimate LD correlations, ∼70% prediction accuracy was preserved in SBayesRC, whereas neither LDpred2 nor SBayesR was able to reach to convergence. For the scenario of unequal per-SNP sample sizes, we simulated two cohorts with a proportion of SNPs genotyped in only one cohort (i.e., non-overlapped SNPs) and obtain the summary statistics from the meta-analysis (**Methods**). As the proportion of overlapped SNPs decreased, SBayesR more often failed in convergence and LDpred2 had a faster rate of decrease in prediction accuracy than SBayesRC (**Fig. 2b**). Of note, LDpred2 tended to preserve prediction performance at the cost of biased estimates in SNP-based heritability and polygenicity (**Supplementary Fig. 6**). In contrast, in SBayesRC, the impact of model misspecification was mostly absorbed in the residual variance which is a nuisance parameter in the model, with no or only marginal bias to the other important genetic architecture parameters (**Supplementary Fig. 5-6**). These results demonstrated that SBayesRC is more robust than the competing methods to the model misspecifications that are likely to occur in practice.

**Figure 2.**
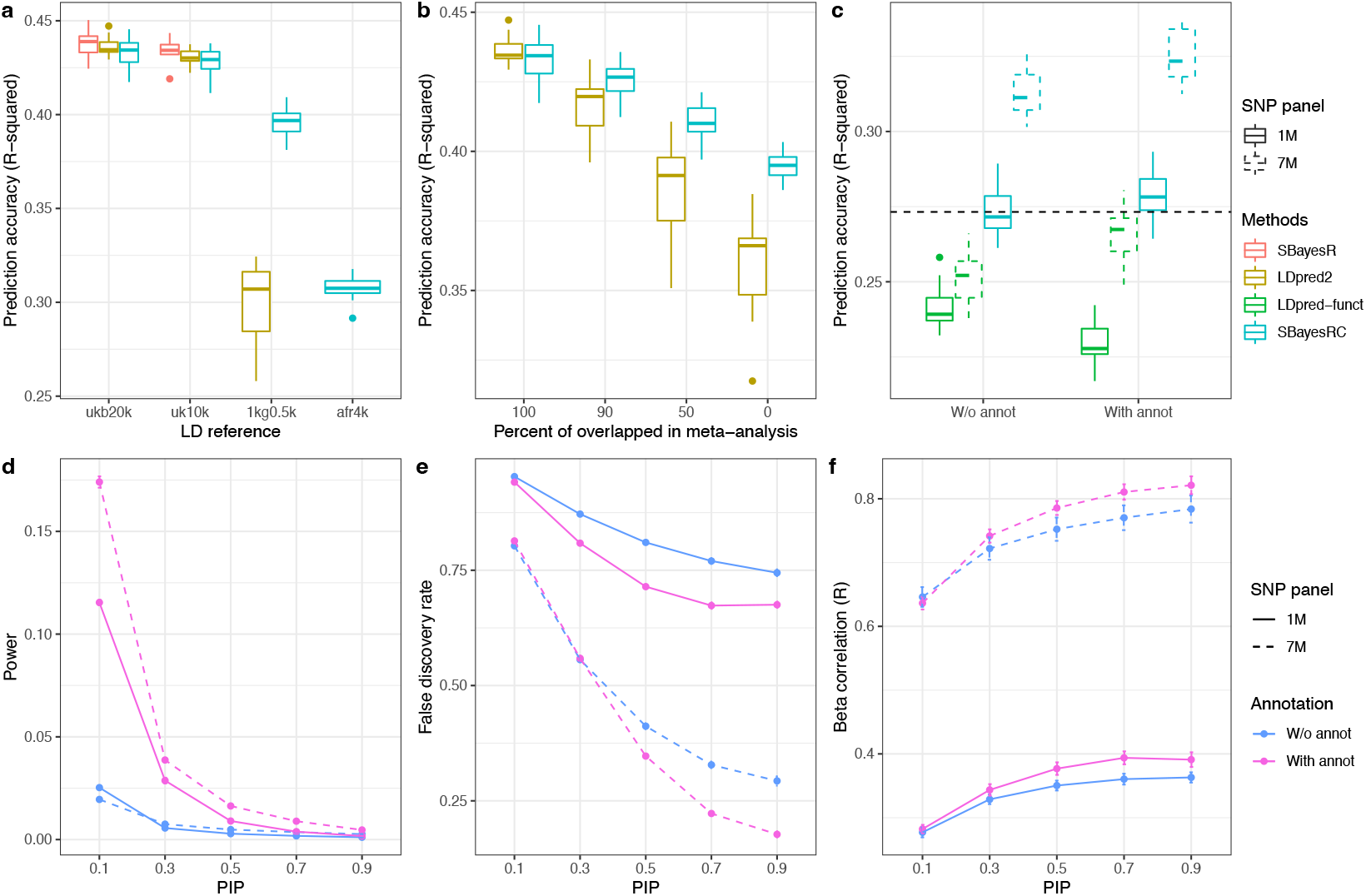
Assessing the performance of our methods by simulations. a) Robustness of SBayesRC to the choice of LD reference. b) Robustness of SBayesRC to the unequal per-SNP sample sizes in the meta-analysis. c) Improvement of prediction accuracy using SBayesRC with high-density SNPs, annotation data, or both, in comparison to LDpred-funct. d) Power of identifying causal variants using models with or without high-density SNPs or annotation data. e) False discovery rate (FDR) of identifying causal variants using model with or without high-density SNPs or annotation data. f) Correlations between the estimated and true effect sizes at SNPs with posterior inclusion probability (PIP) greater than a threshold.

We next evaluated the benefit of using functional annotation data and considering more SNPs beyond the HapMap3 panel (a commonly used panel in the current prediction methods such as LDpred2 and SBayesR). We expanded the simulation to include 7,356,518 imputed common SNPs and incorporated functional annotations to simulate the causal effects. In the simulation, we sampled the causal variants from the 7M SNPs, with their effects sampled from a mixture distribution depending on the observed functional annotations from the curated annotation dataset^24^ (LD score regression baseline model BaselineLD v2.2) and the annotation effects estimated from the SBayesRC analysis in UKB height (**Methods**). As expected, we observed a significant improvement in prediction accuracy when using more SNPs and/or annotation data in SBayesRC (**Fig. 2c**). Compared to using 1M HapMap3 SNPs, using all 7M SNPs improved the prediction accuracy by 14.4% (the difference in prediction 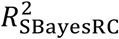 between 7M and 1M SNPs divided by that with 1M SNPs). Compared to the no annotation model, the model incorporating annotation data improved the prediction accuracy by 2.0% and 3.8% when using 1M HapMap3 and 7M common SNPs, respectively. A similar pattern was observed in LDpred-funct that using annotation data on 7M SNPs gave higher prediction accuracy than that on 1M SNPs, but the prediction accuracy was substantially lower than that from our method in each scenario (**Fig. 2c** and **Supplementary Fig. 7**). We hypothesize that the advantage of exploiting annotations is because of both better identification of causal variants and better estimation of their effect sizes. This hypothesis is supported by the results that for a given PIP threshold, incorporating annotations in the model led to a higher power and a lower false discovery rate (FDR) for identifying the causal variants (**Fig. 2d,e**) and a stronger correlation in the estimated and true SNP effects (**Fig. 2f**). Coupled with the higher prediction accuracy, the SNP-based heritability estimate increased toward to the true value in the simulation when more SNPs with annotation data were used (**Supplementary Fig. 9**).

### Improved prediction accuracy within European ancestry

We performed 10-fold cross-validation in the UKB sample and cross-biobank prediction in the Lifeline cohort to assess the performance of SBayesRC for predicting individuals of European ancestry (**Methods**). As in the simulation, the per-SNP annotation data were from BaselineLD v2.2^24^, which included 83 binary annotations such as coding, promoter, enhancer, or conserved variants and 13 quantitative annotations such as predicted allele age and background selection statistic. There were in total 10M imputed common SNPs in the UKB. After matching SNPs across GWAS, validation and annotation data sets, 7M common SNPs were retained for the subsequent analyses (**Methods**).

In the UKB cross-validation, we analysed 28 independent traits including 8 diseases (**Supplementary Table 1**). We compared the performance of our method using 7M SNPs, annotation data, or both, to that of common practice of using 1M HapMap3 SNPs without any annotation, and to the performance of other methods including C+PT^44^, SBayesR^35^, LDpred2^43^ and LDpred-funct^31^. Other than C+PT, all these methods are joint-effect models that generate joint SNP effects as predictors, in which SBayesR and LDpred2 are widely used methods for analysing HapMap3 SNPs and LDpred-funct is the latest method that can analyse 7M SNPs and incorporate annotations (**Table 1**). For the ease of comparison, the prediction accuracy from each method was calculated as the relative value to that from SBayesR from out-of-sample prediction, i.e., relative prediction accuracy of method 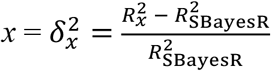, where *R*^2^ is the prediction accuracy. We report the mean 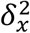 over 10-fold cross-validation. When using HapMap3 SNPs only, SBayesRC without annotations gave approximately the same prediction accuracy as SBayesR, which was significantly higher than that of LDpred2 (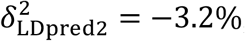, Wilcoxon signed rank exact test *P*-value = 1.4 ×10^−7^) (**Fig. 3a**). Compared to using HapMap3 SNPs without annotations, the prediction accuracy was improved by 2.8% (*P*=0.001) or 3.2% (*P*=3.2 × 10^−7^) corresponding to using either 7M SNPs or annotation data in our method. When using both 7M SNPs and annotation data, the relative prediction accuracy increased to 14.2% (*P*=7.5 × 10^−9^), substantially higher than using either source of information alone, suggesting a strong interaction between the SNP density and annotations (see more discussion below). Although more SNPs together with annotation data also improved the mean relative prediction accuracy from LDpred-funct, it was not significantly greater than that of the baseline method (max 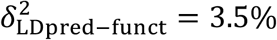; *P*=0.58), i.e., SBayesR using 1M HapMap3 SNPs without annotations. In LDpred-funct, results from that traits varicose veins, risk taking and waist-hip ratio (adjusted for BMI) were large outliers which all had a relatively low SNP-based heritability 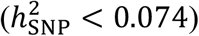 and low baseline SBayesR R^2^ 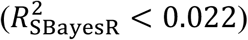; in addition, although the prediction accuracies for these traits were relatively high, the regression slopes were largely biased (**Supplementary Table 2-3**). On average across all traits, SBayesRC gave 11.9% higher prediction accuracy than LDpred-funct using 7M SNPs and annotation data (*P*=5.5 × 10^−5^). In line with the literature^21,45-47^, all of the above joint-effect models were markedly superior to the C+PT approach (a marginal-effect model; 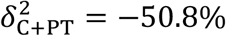 *P*=7.5 × 10^−9^), which did not show any improvement from using more SNPs, consistent with the finding of a recent study^48^. In addition to high prediction accuracy, SBayesRC gave regression slopes close to one across different traits, suggesting no significant prediction bias, better than those from other methods (**Supplementary Fig. 12**).

**Figure 3.**
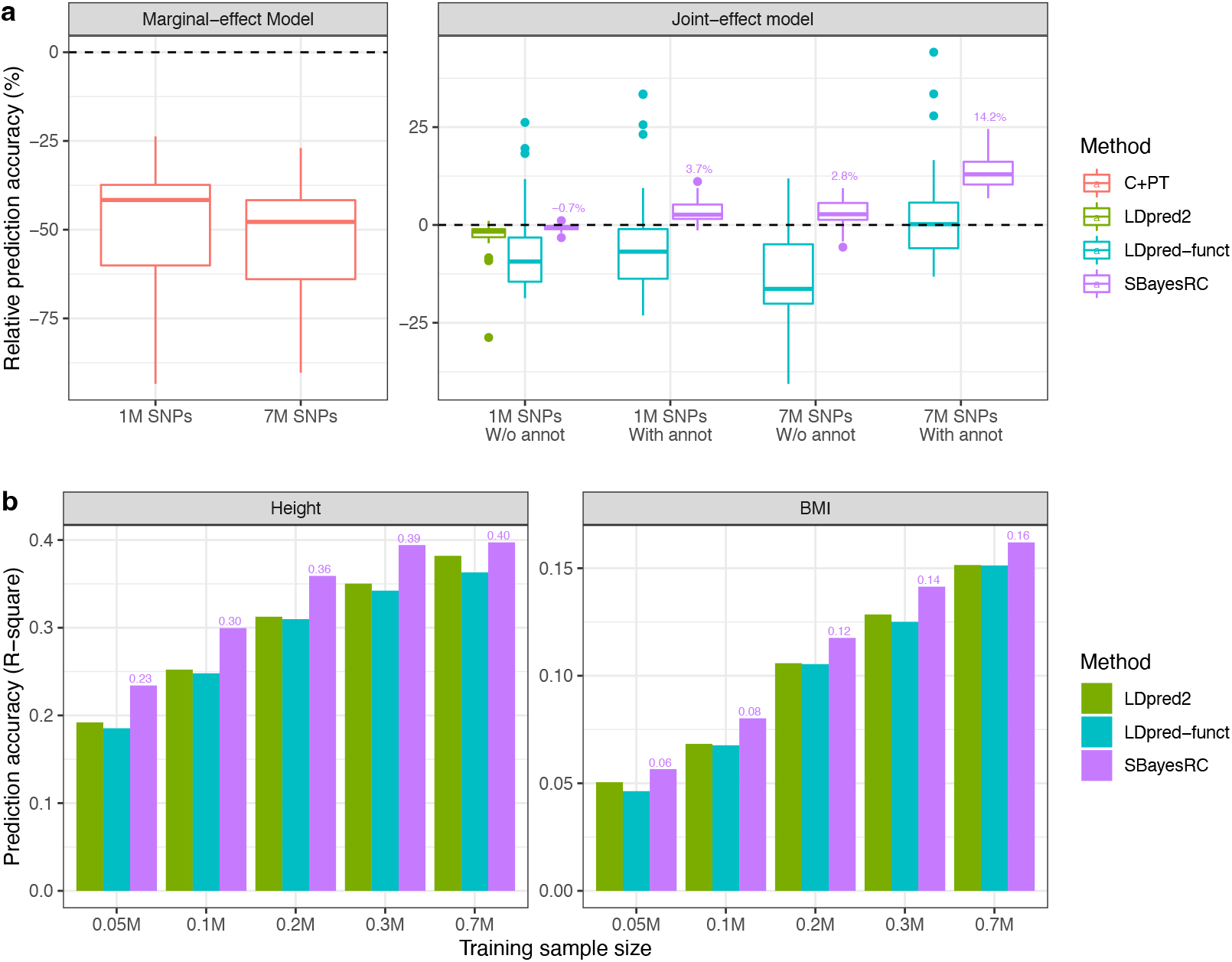
Prediction performance using SBayesRC with 7M SNPs and annotation data in European populations. a) Relative prediction accuracy of different methods to SBayesR using 1M HapMap3 SNPs, averaged from ten-fold cross-validation in the UKB. b) Out-of-sample prediction accuracy for height and BMI, using UKB (N = 0.05 to 0.3M by down sampling) or GIANT dataset^49^ (N = 0.7) as training and an independent dataset of Lifeline as validation.

In the cross-biobank prediction analysis, we focused on the analysis of height and body mass index (BMI) for which GWAS summary statistics are publicly available with different sample sizes. We used summary statistics from UKB and GIANT consortium^49^ (sample size up to 0.7 million) for training, and validated in the Lifeline cohort (n=11,842)^41^. All the training and validation datasets were of European ancestry. As expected, the prediction accuracy improved as the training sample size increased for both height and BMI in all methods. SBayesRC had the highest prediction accuracy in each sample size, outperforming LDpred2 by 4.0-21.9% (which only used 1M HapMap3 SNPs without annotations), and outperforming LDpred-funct by 7.1-26.3% (which used the same data as SBayesRC but required additional tuning dataset) (**Fig. 3b**). Using the largest sample size to date (n_GIANT_=0.7M) and SBayesRC with 7M SNPs and 96 per-SNP annotations, we achieved a maximum prediction *R*^2^ of 0.40 for height and 0.16 for BMI in the Lifeline cohort, explaining 65% and 73% of SNP-based heritability estimated by SBayesRC (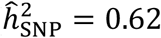for height and 0.22 for BMI), which serves as the theoretical upper bound of the out-of-sample prediction accuracy for each trait.

### Improved accuracy in trans-ancestry prediction

To test whether the superiority of using functional annotations with genome coverage for prediction is transferable into other populations of different ancestries, we performed trans-ancestry prediction in UKB, where predictors derived from GWAS in European ancestry were validated in samples of South Asian (SAS), East Asian (EAS), and African (AFR) ancestries, respectively. In this analysis, we calculated two types of relative prediction accuracy for each trait. In the first type, the benchmark was the prediction accuracy of SBayesR with 1M HapMap3 SNPs averaged across 10 folds of cross-validation in the European ancestry. In the second type, the benchmark was the prediction accuracy of SBayesR trained in the GWAS of European ancestry and validated in each of the other ancestries. To reduce sampling variation in relative prediction accuracy, we removed 10 traits with prediction accuracy (R^2^) from SBayesR less than 0.05 in the cross-validation in European ancestry and one trait failed to converge with PolyPred-S (age at menarche), retaining 17 traits for the trans-ancestry prediction.

We compared our method to two recently developed methods designed for trans-ancestry prediction, PolyPred-S^50^ and PRS-CSx^14^ (**Table 1**). Because both methods require a tuning sample of individual-level data from the target population to generate the final SNP weights for prediction, we held out a sample of 500 individuals from the validation dataset for each non-European ancestry for SNP weights tuning. Given the genetic discrepancies between ancestries, the tuning step is critical and could override the differences between models in themselves. For a fair comparison, we also allowed our approach to use this tuning data by first running SBayesRC in each of the GWAS datasets of different ancestries separately and then combining the SNP effects with weights estimated from the tuning data (we refer this analysis strategy to as SBayesRC-multi; **Methods** and **Table 1**).

Relative to the prediction accuracy within European ancestry, the trans-ancestry prediction accuracy attenuated with increased genetic distance to the European ancestry, regardless of methods (**Fig. 4a**), in line with previous studies^14-19,48^. Despite the overall decline in prediction accuracy, the use of high-density SNPs beyond HapMap3 or annotation data increased prediction accuracy when benchmarking with that of SBayesR observed within each of the ancestries (**Fig. 4b**). Consistent with the patterns observed within European ancestry, SBayesRC using both 7M SNPs and annotation data gave the highest prediction accuracy, with a relative improvement of 16.0% in SAS (*P*=1.5 × 10^−5^), 22.6% in EAS (*P*=2.1 × 10^−4^), and 33.7% in AFR (*P*=4.6 × 10^−5^), averaged across traits. In contrast, we did not observe large improvement using PolyPred-S, except in the AFR population (mean relative prediction accuracy of 2.6% with *P*=7.9 × 10^−3^ in SAS; 2.0% with *P*=0.55 in EAS; 29.9% with *P*=6.7 × 10^−3^ in AFR). On average, SBayesRC outperformed PolyPred-S by 13.2%, 19.7% and 13.2% in SAS, EAS and AFR, respectively (mean improvement of 15.4% across the three non-European ancestries). A notable outlier was vitamin D which had 331% improvement in PolyPred-S on the basis of 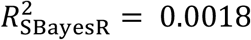 in AFR. This remarkable improvement is likely achieved from the use of tuning sample in PolyPred-S, which can help to identify causal variants that are in high LD in EUR but have very different LD in AFR. Using additional set of GWAS summary statistics from Biobank Japan (BBJ^51^), PRS-CSx outperformed SBayesRC by only 0.4% for predicting EAS individuals in the UKB but was inferior to SBayesRC-multi by 13.5% (**Fig. 4c**). Note that the summary statistics were separately analysed in SBayesRC-multi but were jointly modelled in PRS-CSx. The fact that SBayesRC-multi was favoured suggests that the benefit of leveraging functional annotations with all imputed SNPs outweighs the benefit of joint modelling summary statistics from multiple populations at a subset of SNPs.

In addition to improved prediction accuracy, SBayesRC used less computational resources than other methods (**Table 2**). For the analysis of 7M SNPs with 96 annotations, SBayesRC required 75 GB RAM and 9.5 computing hours per core with 4 CPU cores, which are commonly available in a standard computing cluster.

**Figure 4.**
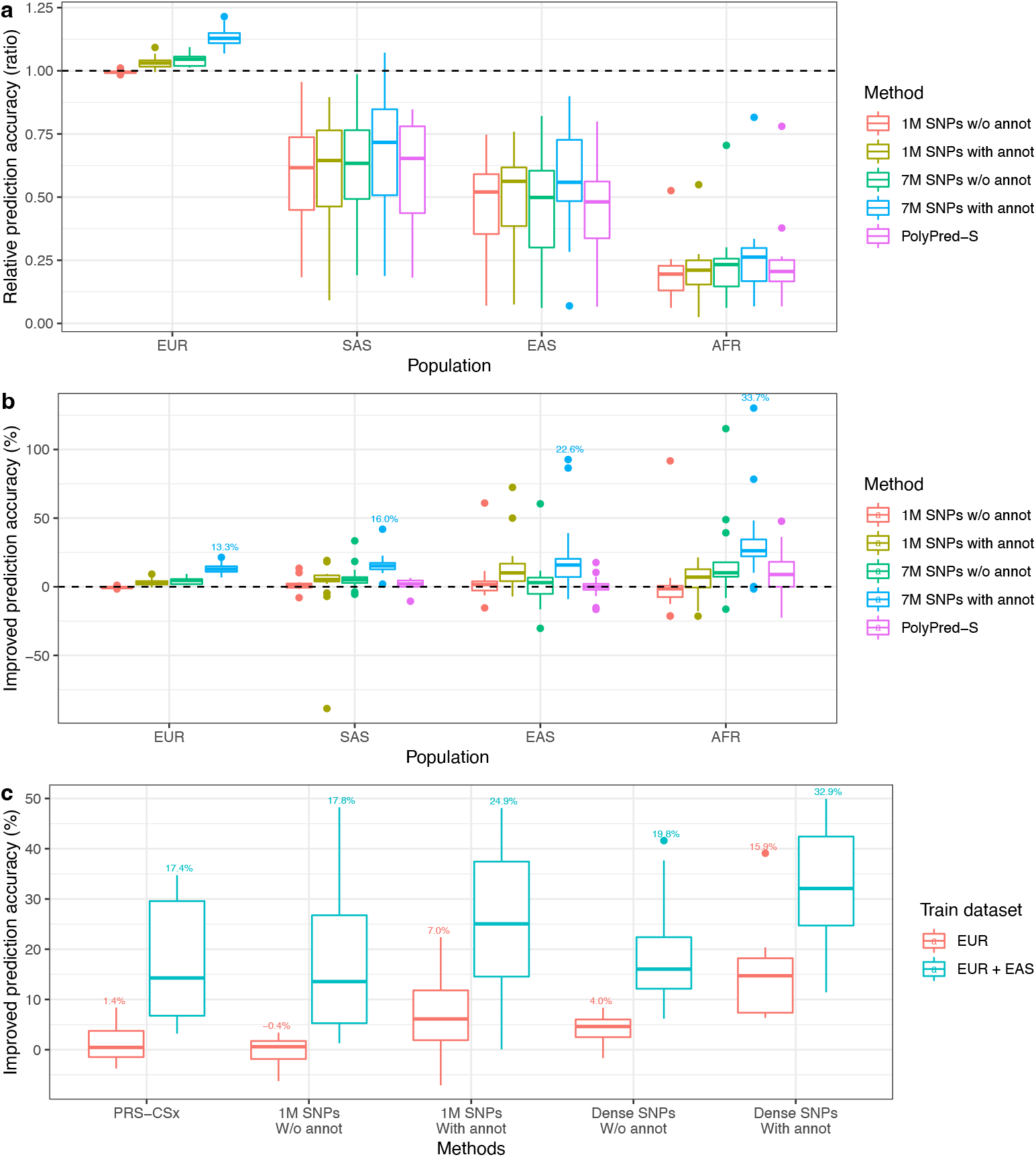
Trans-ancestry prediction using SBayesRC with 7M SNPs and annotation data. a) Relative prediction accuracy (ratio) to that of SBayesR with 1M HapMap3 SNPs averaged across 10 folds of cross-validation in the European ancestry. b) Relative prediction accuracy (% of improvement) to that of SBayesR trained in the GWAS of European ancestry and validated in each of the other ancestries. VitD in PolyPred-S AFR population had a value of 331%, which is removed from the graph for a clear presentation. c) Relative prediction accuracy as in panel b either using summary statistics from UKB of European ancestry alone or together with those from BBJ of East Asian ancestry for trans-ancestry prediction in the UKB population of East Asian ancestry (8 traits available). The number above each boxplot indicates the mean value across traits. Data provided in **Supplementary Table 4 and 5**.

### Significant interaction between SNP density and annotation information

Results above have shown that using a full imputation SNP set (i.e., not limiting to the HapMap3 panel) or using annotation data can each lead to a significant improvement in prediction accuracy. Moreover, using 7M SNPs and annotation data together remarkably outperformed using either one source of information alone, suggesting an interaction effect between SNP density and annotation information for prediction. To test and quantify the significance of this interaction effect, we calculated the relative prediction accuracy of exploiting annotations in the model, i.e., 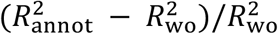 Where 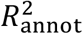 is the prediction accuracy with annotations and 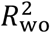 is that without annotations, given a set of SNPs. As shown in **Fig. 5**, using 7M SNPs resulted in a significantly higher relative prediction accuracy of exploiting annotations than using 1M HapMap3 SNPs in both simulation and real trait analyses, manifesting a significant interaction effect. More specifically, regardless of the level of heritability in simulation, sample size of GWAS in real traits, within- or cross-ancestry prediction, the use of 7M SNP annotations (y) led to about twice as much improvement as that of using 1M SNP annotations (x) (**Fig. 5**; regression slope = 1.88 with *P*=1.4 × 10^−5^ from regression of y on x weighted by the number of traits in each data point). For example, in the cross-validation of complex traits in the UKB European sample, incorporating annotations increased prediction accuracy by 4.4% with 1M HapMap3 SNPs but by 11.2% with 7M imputed SNPs. This is consistent with our hypothesis that the annotations at the SNPs in LD with a causal variant may not track the annotation at the causal variant, resulting in a loss in information (**Fig. 1a**). To pursue the maximum improvement, we further considered all imputed common variants in the UKB (∼10M SNPs) by extracting their annotations from the PolyPred model (**Methods**), but did not observe more improvement in prediction accuracy (**Supplementary Fig. 15**), suggesting a saturation of association signals detectable by imputation. Thus, it is likely that the benefits of leveraging annotation data for polygenic prediction will be maximized when analysing variants at sequence level.

**Figure 5.**
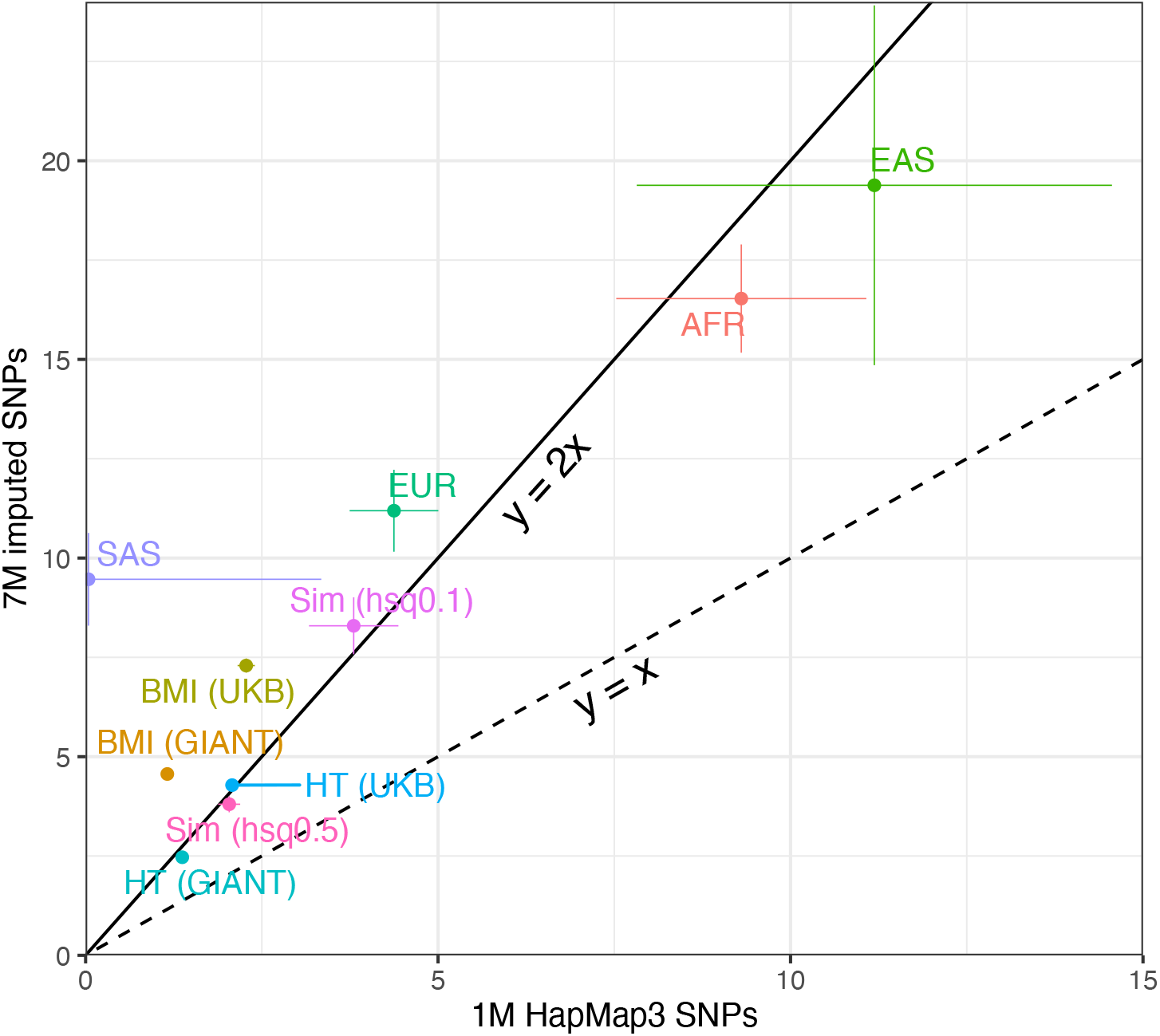
Comparison between 1M HapMap3 SNPs and 7M imputed SNPs for the improvement (%) in prediction accuracy for SBayesRC using annotations relative to SBayesRC without annotations. The slope of regression of results from 7M SNPs on results from 1M SNPs is 1.88 (s.e.=0.22).

### Other factors affecting accuracy of prediction leveraging functional annotation

Here we investigate other factors, besides SNP density, that affect accuracy of prediction leveraging functional annotations, including SNP-based heritability, GWAS sample size, properties of minor allele frequency (MAF) and LD, the number of annotations, and the strategy of analysis.

Stratifying traits based on the SNP-based heritability estimates from SBayesR found that traits with lower SNP-based heritability tended to benefit more from exploiting annotation data on 7M SNPs (regression slope = -15.7, *P*=0.031; **Fig. 6a**). When down sampling the UKB data for GWAS in height and BMI, the relative prediction accuracy using 7M SNPs and annotations was higher with a smaller GWAS sample size, whereas using more SNPs alone required a larger sample size to give a higher improvement in prediction accuracy (**Fig. 6b**). This is expected because including annotations in the model is analogous to adding more data as annotations are independent of genome variations, whereas including more SNPs in the model alone increases the number of parameters to estimate therefore consuming more degrees of freedom. Given that most traits have a polygenic architecture, low SNP-based heritability or small sample size means low power in GWAS. Thus, results from this analysis suggest that traits with insufficient power would benefit more from leveraging annotation data for prediction.

**Figure 6.**
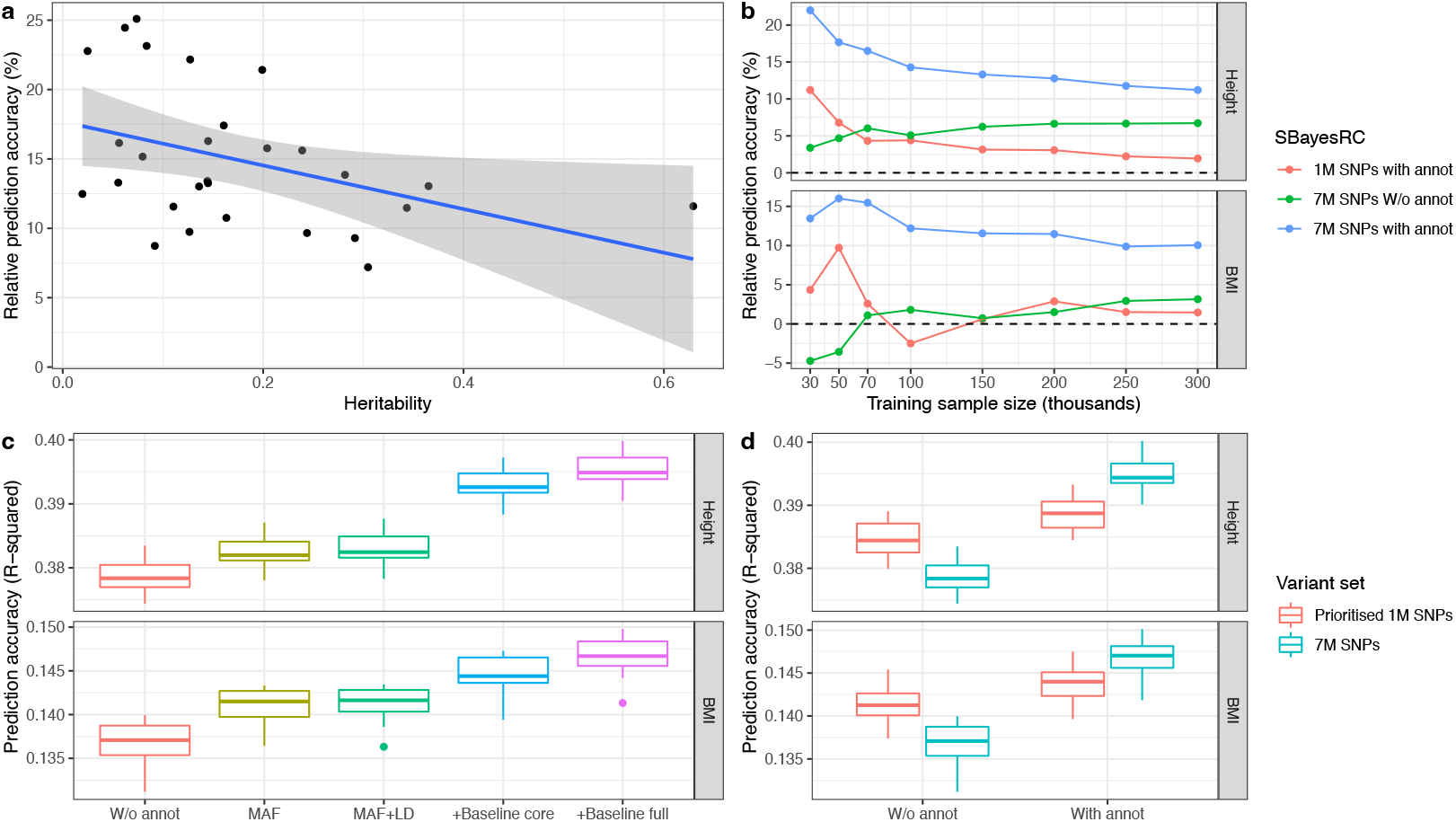
Other factors affecting accuracy of prediction incorporating functional annotations.a) Traits with low heritability tend to benefit more from using annotation data. b) GWAS with small sample sizes tend to benefit more from using annotation data. c) Improvement in prediction accuracy increases with the number of annotations upon the MAF and LD (+ Baseline core/full = MAF+LD+Baseline core/full). d) Full analysis of all SNPs and annotation data is superior to the stepwise analysis that prioritises the top 1M SNPs based on their annotations and fits them in the model.

To investigate whether functional annotations provide additional information than LD and MAF properties, we evaluated the relative prediction accuracy for height and BMI by first fitting 13 MAF related annotations from BaselineLD only and gradually adding 4 LD bins, 21 core annotations^52^, and finally the full set of 96 annotations. We found that the relative prediction accuracy increased with just fitting MAF, had no significant change when adding LD bins, and further increased with additional functional annotations (**Fig. 6c**). In addition, conditional on MAF and LD bins, the improvement was larger with 96 functional annotations than that with 21 core annotations. These results suggest that functional annotations are more informative than LD and MAF annotations, and using a rich set of functional annotations altogether is better than using a few key functional categories.

Lastly, we compared alternative strategies of analysing all common SNPs with annotations. Simultaneous fitting of 7M SNPs in the model is computationally impractical in most MCMC-based methods. One strategy is to perform a stepwise analysis^53,54^ as below. Step 1, estimation of impact of functional annotations on SNP effects, which is often quantified by estimation of the annotation-specific enrichment in per-SNP heritability using S-LDSC^38^. Step 2, prioritisation of all SNPs based on their functional annotations and results in step 1. Final analysis uses a subset of the SNPs that rank from the top in step 2. To compare our method to such a stepwise strategy, we selected the top 1M SNPs from 7M SNP set based on the results from S-LDSC^38^, and performed the prediction analysis in height and BMI. For SBayesRC in which annotations were not assigned, the functionally-prioritised 1M SNP set had a higher prediction accuracy than the 7M SNPs, whereas for SBayesRC in which annotations were assigned, the functionally prioritised 1M SNPs was slightly better than the same set of SNPs without re-estimation the annotation effects but was inferior to the 7M SNPs (**Fig. 6d**), suggesting that the unified analysis is better than the stepwise analysis in refining the information from annotation data.

### Contributions of functional categories to prediction accuracy

We have shown that functional annotations are useful to improve polygenic prediction. We then sought to identify which functional annotations are of most importance. To this end, we constructed functional category-specific PGS using SNPs within that functional category and their effect estimates from the genome-wide analysis of SBayesRC. Overall, categories with more SNPs contributed more to the prediction accuracy, but with apparent outliers (**Fig. 7a**). We found that evolutionary constrained regions, albeit small in SNP set size, had the greatest contribution among all categories without flanking windows. For example, regions that are conserved across 29 eutherian mammals (Conserved_LindbladToh^55^ in BaselineLD) only covers 2.9% of the genome but contributed 40.5% of prediction accuracy averaged across traits, resulting in 14.0-fold enrichment in the per-SNP contribution to prediction accuracy (per-SNP predictability enrichment = 40.5/2.9). For comparison, the coding regions (1.6% of the genome) contributed 25.9% of prediction accuracy, with a per-SNP predictability enrichment fold of 16.5. This result suggests that evolutionary constrained variants are as informative as the coding variants with respect to complex trait prediction, despite only 33.2% coding variants are annotated as constrained variants. Across functional categories, the per-SNP contribution to prediction accuracy was in proportion to the per-SNP contribution to heritability (**Fig. 7b**), suggesting that the variance explained by a SNP in the GWAS sample can be transferred into predictive ability of the SNP in the validation sample. The largest per-SNP predictability was from the non-synonymous SNPs in the coding sequence (41.4-fold enrichment), which also had the largest enrichment in per-SNP heritability.

**Figure 7.**
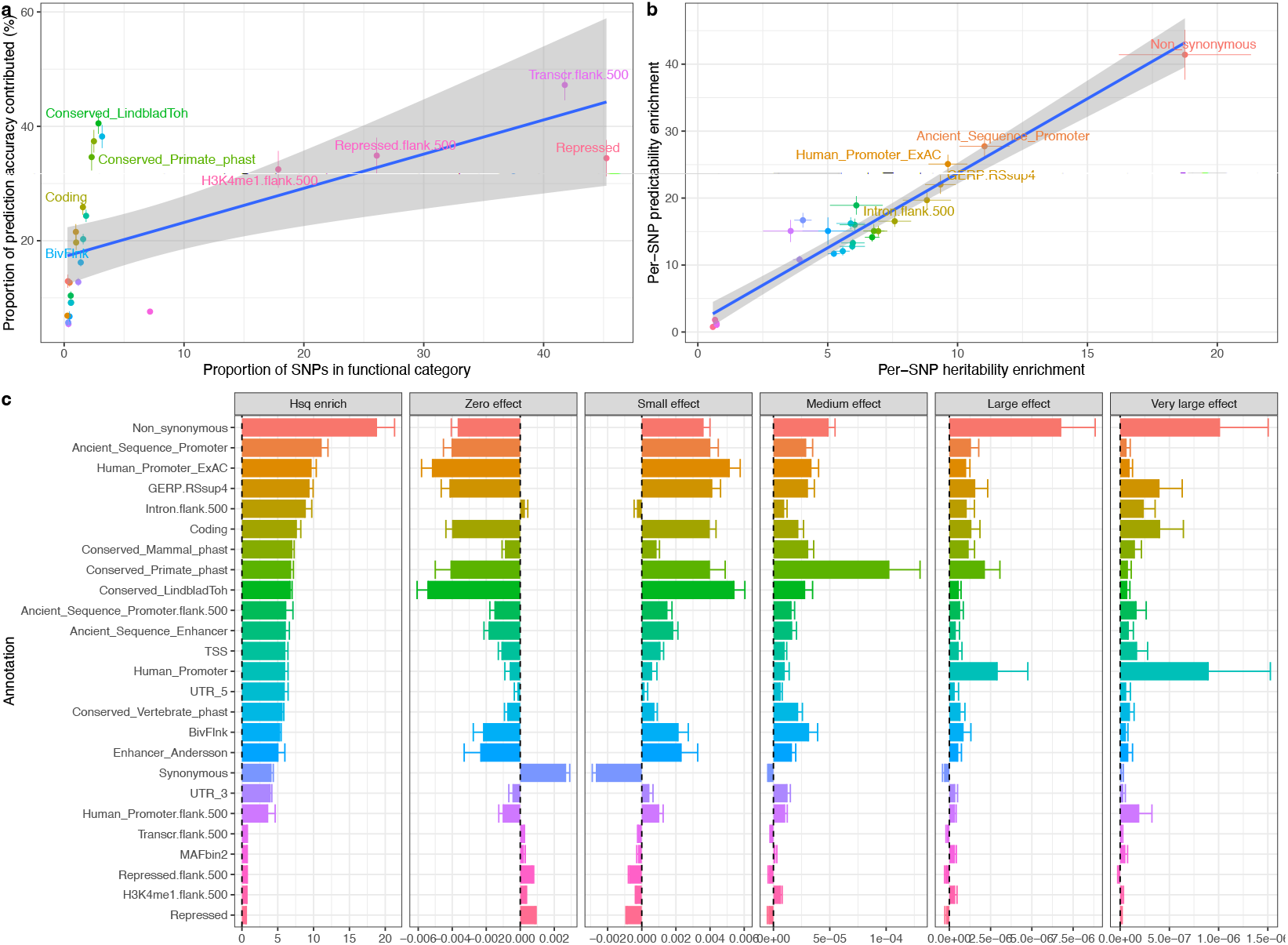
Contribution of functional categories to the total prediction accuracy and estimation of functional genetic architecture in complex traits. a) Proportion of prediction accuracy against proportion of SNPs in each functional category. b) Per-SNP contribution to prediction accuracy against per-SNP contribution to heritability in each functional category. c) Per-SNP heritability enrichment and distribution of effect sizes shown by the proportion of SNP effects belonging to each of the five mixture distributions across the top 20 and bottom 5 functional categories.

We prioritised functional annotations based on their per-SNP heritability enrichment averaged across traits analysed in this study. The top 20 had mean fold enrichment in per-SNP heritability, which ranged from 3.8 to 18.8, included non-synonymous variants, evolutionary constrained regions, coding sequence and regulatory elements, by-and-large consistent with the results from S-LDSC (**Supplementary Fig. 16**). Our method allows us to go on ask whether the enrichment in per-SNP heritability is due to more causal variants or larger effect sizes in the category. We found that conditional on the other annotations, the non-synonymous SNPs category were enriched in both the proportion of causal variants and the magnitude of effect sizes (**Fig. 7c**). Moreover, compared to evolutionary conserved regions in mammals, conserved regions in primates had lower proportion of null SNPs and higher proportions of SNPs with small to large effects in human traits.

## Discussion

We have introduced a novel method, SBayesRC, for polygenic prediction of complex traits using GWAS summary statistics of the full set of imputed SNPs and incorporating diverse functional annotations on each SNP. Compared to the common practice of using 1M HapMap3 SNPs, using 7M imputed common SNPs and 96 per-SNP annotations, we observed 14% improvement in prediction accuracy within European ancestry averaged over 28 complex traits and diseases, and up to 33% across ancestries averaged over 18 well powered traits. Our observation of improved prediction exploiting annotation data is consistent the findings in the previous studies^15,29,30,33,46,54^. Our method outperformed the best method for within European ancestry prediction using annotations, LDpred-funct, by 12%, and the best method for cross-ancestry prediction using annotations, PolyPred-S, by 15% averaged across ancestries (20% in EAS), and that without annotations but jointly models GWAS summary statistics from multiple populations, PRS-CSx, by 14% when the data from the EAS ancestry was also used in our method. Furthermore, we found a significant interaction between SNP density and annotation information for prediction accuracy by observing consistent results across datasets that use of 7M SNPs doubled the benefits of incorporating annotations into prediction as compared to 1M HapMap3 SNPs.

The interaction between SNP density and annotation information is likely due to fewer “errors” in the annotation data when more variants are uncovered through imputation. When the causal variants are unobserved, SNPs used in the analysis can capture the causal effects by LD but may be annotated differently from the annotations at the causal variants. In other word, the causal effects are assigned to other variants with “incorrect” annotations. This would dilute the information from each annotation and smooth out the differences in effect size distribution between annotations if the unobserved causal variants are distributed at random, or would cause a systematic bias in the annotation effect estimates if the unobserved causal variants are enriched in some annotation categories. In either case, the annotation data are used less effectively. As shown in the simulation study, the estimation for the proportion of SNPs in each non-zero distribution was biased when using the 1M subset of SNPs (**Supplementary Fig. 10**), in concordance with the lower prediction accuracy than using a full SNP panel including the causal variants (**Supplementary Fig. 11**). In the real data analysis, we observed a significant improvement from 1M to 7M imputed SNPs but no further difference to 10M imputed SNPs (**Supplementary Fig. 15**), likely due to the limitation of imputation accuracy. Thus, it is expected that using variants at sequence level will maximize the efficiency of borrowing information from the annotation data for prediction.

We found that the combination of high-density SNPs and functional annotations provides most benefit to traits with low SNP-based heritability or small GWAS discovery sample sizes through provision of additional information to allele frequency and linkage disequilibrium categories, and is best used in a unified analysis where all parameters are estimated in one model. These results highlight the utility of leveraging functional annotations for predicting disease risk, as most common diseases do not have a high SNP-based heritability and the effective sample sizes are still limited for many diseases. These results support the need for generation of more high-quality functional annotations, because empirically they provide extra information in addition to non-functional dependent annotations, such as MAF and LD (none of the MAF or LD categories were present in the top 20 annotation categories ranked based on the per-SNP contribution to prediction accuracy; **Fig. 7c**). Our method can incorporate many annotations and estimate both annotation effects (binary and/or quantitative) and SNP effects from the data without additional tuning steps. Our results suggest that such a unified computational framework is more desirable than the stepwise approaches commonly used in the previous studies^53,54^. These results are useful to inform the experimental design of leveraging functional annotations for prediction in the future.

We proposed a rank reduction approach to account for correlations between marginal effects such that the joint effects of genome-wide SNPs are fitted to a set of observables with independent residuals. There are several advantages of using this low-rank model. First, it substantially improved the model robustness. We showed that our method is robust to the LD differences between GWAS and reference samples and the per-SNP sample size variation resulted from the meta-analysis between cohorts with different genotyping platforms. This is not only owing to elimination of numerous small eigenvalues/eigenvectors in each LD block which are subject to high sampling variation in LD, but also because independence of transformed summary statistics enables the sampling of block-wise residual variance, which introduces a mechanism to manage the convergence issue due to violation of model assumptions. It has been found that SNP effect sizes would blow up during MCMC when the model fails to converge. In this case, the sampled values of residual variances would be large if the SNP effect sizes tend to blow up, which will in turn shrink them back toward zero, preventing failure in convergence. Second, the low-rank model substantially improved the computational efficiency which allows us to fit a large number of SNPs with only a small fraction of computation resource compared to the original model. In theory, SBayesRC is scalable to fit variants at sequence level via calibrating the required proportion of variance in LD explained by the selected eigenvalues/eigenvectors, making it a powerful tool to analyse the incoming large-scale whole-genome sequence data. Third, this low-rank model depicts a general framework that can be applied with different priors as used in other methods.

We noted several limitations in this study. First, although our method is scalable to analysis of whole-genome sequence data, only imputed common SNPs that were functionally annotated were analyzed due to limitations in availability of whole-genome sequence data when conducting the study. We investigated use of up to 10 million imputed SNPs with MAF > 0.01 but did not observe a significant improvement comparing to the 7 million SNP set, suggesting a likely saturation of information from imputation and warranting a follow-up study with sequence variants. Second, our low-rank model requires eigen-decomposition on the LD matrices which can be expensive if they are computed one by one. We performed eigen-decomposition on the 7 million SNP set using parallel computing and provide the results online for use by the research community, assuming that the SNPs in their GWAS summary data match with the SNPs we used to generate the LD data. This could be an issue if some SNPs in the LD data are not included in the GWAS, although most studies impute to the same reference data we used (HRC reference panel^56^). One approach to address this is to impute the summary statistics for those “missing” SNPs^57^ (**Supplementary Note**). We found empirically that the loss of prediction accuracy was marginal unless the missing rate was greater than 40%. Third, our method is robust to LD and per-SNP sample size variation, which has been found to be the major cause of the convergence problem in SBayesR and other summary-statistics based methods, but it is still subject to errors in the GWAS summary statistics, such as genotyping errors and allelic mislabeling. Thus, application of additional quality control on the summary statistics prior to the analysis may be required in some circumstances^58^. Fourth, for trans-ancestry prediction, there is possibly further improvement in prediction accuracy by modelling summary statistics from multiple populations jointly as in PRS-CSx^14^, but we leave such an extension of our method to a future project. Fifth, in theory, our method can be applied to a very large number of variants given a memory usage by calibrating the rank reduction parameter, but a too low threshold value may result in a loss of prediction accuracy and therefore requires calibration with caution. Sixth, this study used general annotations curated by the BaselineLD model^38^, which does not include annotations from recent studies using single-cell sequencing technology that uncover more cell-type specific epigenetic marks and chromatin states^59-62^. Given mounting knowledge about which tissues or cell types are relevant to complex traits inferred from GWAS data and single-cell RNA-seq data, using annotations derived from the trait-relevant tissues or cell types are expected to generate more accurate predictors.

In conclusion, the method proposed in this study is a powerful tool to improve polygenic prediction in complex traits and diseases. Our findings provide a guideline how to make best use of functional annotation data for prediction and which functional categories are most useful for within European and trans-ancestry prediction. We anticipate further improved prediction accuracy in the future when the method is applied to whole-genome sequence data with high-quality trait-relevant annotations.

## Methods

### Summary-data-based low-rank model

Consider a general form of the summary-data-based model for fitting SNP joint effects:

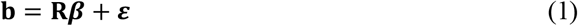

where **b** is the vector of GWAS marginal effect estimates (assuming the genotype matrix **X** has already been standardised with mean zero and variance one), 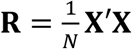 is the LD correlation matrix, *N* is the GWAS sample size, ***β*** is the vector of SNP joint effects, and ***ε*** is the vector of residual terms with 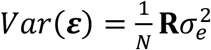. When the marginal effects are estimated from GWAS using genotypes at 0/1/2 scale (**b**^**\***^), **b** can be estimated using **b**^**\***^, standard error, and GWAS sample size (**Supplementary Note**).

Sparse LD matrices estimated from a reference sample are often used to improve computational feasibility, including banded^43,54^, shrunk^35,63^ and block-diagonal^14,64^ matrices. To derive our low-rank model, we use a block-diagonal LD matrix based on the quasi-independent LD blocks found in the human genome^42^. For the best performance, we merge small contiguous blocks to a single block with the minimum width of 4 cM, resulting in 591 merged blocks for the European ancestry. For each block *i*, we perform eigen-decomposition on **R**_*i*_ (the subscript is ignored for simplicity in notation)

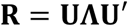

where **U** is the matrix of eigenvectors and **Λ** is the diagonal matrix of eigenvalues. By multiplying both sides of Eq (1) by 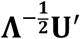, we have

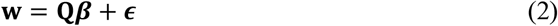

where 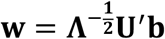 is a linear combination of marginal SNP effect estimates, 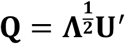 is the new coefficient matrix, and the new residuals 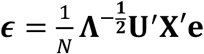 are independently and identically distributed, i.e.,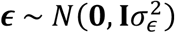. To account for high LD between SNPs and LD variations between GWAS and LD reference samples, we opt to include eigenvectors and eigenvalues for the top principal components (PCs) that collectively explain at least *ρ* proportion of the variance in LD. Assuming *q* top PCs are selected given a value of *ρ*, the dimension of **w** and **Q** is *q* × 1 and *q* × *m*, respectively, with *m* being the number of SNPs in the block. Because *q* is often much smaller than *m*, Eq (2) is a low-rank model and computationally more efficient than Eq (1). We investigated the impact of *ρ* on the method and decided to use *ρ* = 99.5% in practice, which led to a reduction of about 80% rows from the original summary-data-based model at a negligible loss in predictive performance (**Supplementary Fig. 2-3**). In addition, Eq (2) has independent residuals so that it is straightforward to estimate the residual variance, which helps to improve the robustness of the method (**Supplementary Note**). This low-rank model is general and can be applied to any other methods that make different assumptions on the distribution of ***β***. A more detailed derivation is given in the **Supplementary Note**.

### SBayesRC

SBayesRC is a Bayesian method built on the low-rank model described above, assuming a multi-normal mixture distribution for SNP effects. Specifically, we assume that

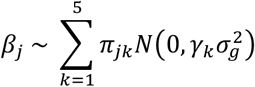

where 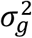 is the total SNP-based genetic variance estimated from the data, ***γ*** = [0, 0.001, 0.01, 0.1, 1]′% depict the scaling factor of five distributions as the mixture components including a distribution of zeros and four normal distributions where each SNP *a priori* explains 0.001% to 1% of genetic variance, and *γ*_*jk*_ is the probability for the SNP effect belong to the *k*^th^ distribution.

In contrast to SBayesR^35^ which assumes the same *γ*_*k*_ for all SNPs, here the probability of distribution membership *γ*_*jk*_ is SNP specific and depends on the annotations on each SNP. Let **A** be the matrix of annotations with a dimension of the number of SNPs *m* by the number of annotations *c*. For each SNP, we model

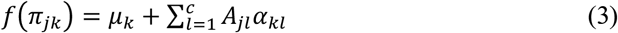

where *f* (·) is a link function that maps the probability variable *γ*_*jk*_ to the real line, *μ*_*k*_ is the intercept capturing the overall proportion of SNPs belong to the *k*^th^ distribution in the genome, *A*_*jl*_ is the value of annotation *l* on SNP *j* (0 or 1 for binary annotations or standardised value with mean 0 and variance 1 for quantitative annotations), and *α*_*kl*_ is the effect of annotation *l* on the membership probability to the *k*^th^ distribution. This generalized linear model allows functional annotations to affect the probability of a SNP being causal (1 – *γ*_*j*1_) and accommodates any distribution of the causal effect (by mixture of a finite number of normal distributions) given the cumulation of functional annotations, regardless of discrete or quantitative annotations, and accounts for overlapping between annotations.

Through estimation of *α*_*kl*_ from the data, this computational framework provides a machinery to make inference on the functional genetic architecture of the trait, because *f* ^− 1^(*α*_*kl*_) quantifies the deviation of the *k*^th^ distribution membership probability, driven by annotation *l*, to the baseline model where all annotation values equal to zero, conditional on the presence of the other annotations. The estimates of *α*_1*l*_, …, *α*_5*l*_ altogether provide a more detailed description about functional architecture than the per-SNP heritability enrichment estimate for an annotation category (**Supplementary Note** and **Supplementary Fig. 10**,**14**). We assume a flat prior for *μ*_*k*_ and a normal prior for 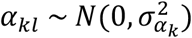 with 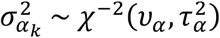 where *υ* _*α*_ = 4 and 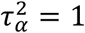.

For a mixture distribution of five components, there are 5 × (*c* + 1) annotation parameters to estimate from the data (including the intercept). In addition, *γ*_*jk*_ is subject to a constraint that 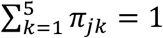 for any SNP. As a consequence, the sampling scheme for *α*_*kl*_ is not straightforward. Although the Metropolis-Hastings algorithm can be used to sample all ***α*** jointly to account for the dependence between elements of ***π***_*j*_, finding the optimal tuning parameters for the proposal distribution could be difficult and specific to trait. To remove the dependence between probability parameters, we employed an alternative parameterization for modelling membership probabilities and annotation effects. Let *δ*_*j*_ be the indicator for the mixture component membership for SNP *j*:

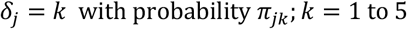

We define a conditional probability that the SNP effect belongs to the *k*^th^ distribution given that it has passed the bar for the (*k*-1)^th^ distribution as

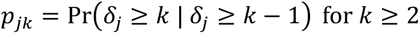

such that *γ*_*j*1_ = 1 – *p*_*j*2_, *γ*_*j*2_ = (1 – *p*_j3_)*p*_*j*2_, *γ*_*j*3_ = (1 – *p*_*j*4_)*p*_*j*3_ *p*_*j*4_, *γ*_*j*4_ = (1 – *p*_*j*5_)*p*_*j*4_*p*_*j*3_*p*_*j*2_, and *γ*_*j*5_ = *p*_*j*5_*p*_*j*4_*p*_*j*3_*p*_*j*2_. We then apply the generalized linear model, Eq (3), to link *p*_*jk*_ with ***α***_*k*_. In this parameterisation, all *p*_*jk*_ are independent, which means that ***α***_*k*_ can be sampled in parallel in each MCMC iteration, and *α*_*kl*_ can be sampled from its full conditional distribution using Gibbs sampling algorithm when the probit link function is chosen, namely *f* (*p*_*jk*_) = Φ(*p*_*jk*_) where Φ(·) is the cumulative density function (CDF) of the standard normal distribution. More details about the alternative parameterization and the Markov chain Monte Carlo (MCMC) sampling scheme are described in the **Supplementary Note**. In all SBayesRC analyses in this study, we ran MCMC for 3,000 iterations with the first 1,000 iterations as burn-in and the rest used for posterior inference. Running a longer chain did not change the prediction accuracy in the simulation and real trait analysis.

### UK Biobank

The UK Biobank (UKB) is a large volunteer cohort with sample size more than 500,000 from across the United Kingdom and has extensive phenotypic and genotypic information from the participants. Informed consent with the protocol’s approval from National Research Ethics Service Committee was signed by all participants. The genotype data was generated using two array chips, the Applied BiosystemsTM UK Biobank AxiomTM Array and the Applied BiosystemsTM UK BiLEVE AxiomTM Array. The UKB analysis team conducted the SNP imputation with reference panels from Haplotype Reference Consortium^56^ (HRC) and the UK10K project^65^. We called the imputed data to BED format by PLINK with best-guest calling, kept the SNPs with MAF ≥ 0.01, Hardy-Weinberg Equilibrium test P ≥ 10^−10^, imputation info score ≥ 0.6 in European samples. We used the GCTA software^66^ to remove the cryptic relatedness in the UKB based on the HapMap3 SNPs in each population. The samples were pruned by estimated relatedness larger than 0.05, keeping the unrelated samples only. We also removed the samples with mismatched sex information in phenotype and genotype, and samples withdrew the participation. The final dataset contains 4 ancestries, European (EUR, N= 347,800), East Asian (EAS, N=2,252), South Asian (SAS, N=9,436) and African (AFR, N=7,006).

We matched the SNPs between UKB, the annotation baseline model BaselineLD v2.2 and LifeLine cohort, extracted the common SNPs among those 3 datasets, resulting in 7,356,518 SNPs and 1,154,522 HapMap3 SNPs. For a secondary analysis, we further included up to 9,705,522 imputed common SNPs with their annotation data extracted from PolyPred-S, which used BaseLineLF (an extended version of BaseLineLD v2.2 to include annotations at the low-frequency variants). We randomly sampled 5,991 EUR samples as the tunning sample for C+PT and LDpred2 and performed 10 cross validation in the remaining samples (341,809). The 55 traits with relatively large sample size (*N* > 110,000) were extracted from all 4 ancestries. The phenotypes with continuous values were filtered within the range of mean +/-7SD and then rank-based inverse-normal transformed within each ancestry and sex group.

### 1000 Genomes and UK10K data

In addition to the LD reference from UKB, we also used two other whole genome sequence data sets. We downloaded genotype data from 1000 Genomes Project (phase 3)^67^. We kept samples labelled as “GBR”, “CEU”, “TSI”, “IBS” and “FIN” as European samples. We extracted the same SNP sets QCed above (7,356,518), and removed the cryptic relatedness based on the HapMap3 SNPs by GCTA software (cut-off value for GRM 0.05), resulted in 494 unrelated samples. We used the genotype data from UK10K project^65^ which consists of 3781 individuals and 45.5 million genetics variants. We extracted the SNP sets QCed above and kept the 3642 unrelated samples by GCTA software.

### Lifeline cohort

From the Lifeline cohort we used 36,305 samples and 17 million SNPs after imputation and QC (imputation info score > 0.3, MAF > 0.0001 and HWE > 1e-6). We kept the sample with age > 20 years old and removed the samples with the phenotypic value (height and BMI) out the range of mean +-5SD. We further removed the related samples and retained 11,842 unrelated samples for out-of-sample prediction.

### Public data from GWAS meta-analysis for height and BMI

We assess the prediction accuracy for height and BMI from the summary data public available^49^. We kept the same variant set in the UKB, extracted the SNPs with per-SNP sample size within mean +/-3 SD and the difference in allele frequency between GWAS and LD reference samples smaller than 0.2. The summary data was further QCed by DENTIST^58^ to filter the SNPs with potential errors, removed the SNPs with P_DENTIST < 5 10^−8^ and P_GWAS > 0.01. All the summary data were imputed to the same variant panel.

### Cross-validation in UKB

We performed ten-fold cross-validation in the UKB with 341,809 unrelated individuals of European ancestry. We chose 55 traits from the UKB with relatively large sample size and pruned the traits with pair-wise phenotypic correlation |r| < 0.3. 31 independent traits were selected for the prediction analysis, including 11 binary traits and 20 continuous traits, where 3 binary traits were further removed due to a very low average prediction accuracy (mean R^2^ < 0.01 among all methods in the European cross-validation). The sample sizes for the final set of 28 independent traits are shown in **Supplementary Table 1**. We partitioned the total sample into ten equal-sized disjoint subsamples. For each fold, we retained one subsample as validation and other remaining nine subsamples as training data. We repeated the process ten times. The summary statistics for each fold were generated by PLINK2 software^68^ with sex, age, and first 10 principal component as covariates. Linear regression was used for continuous traits and logistic regression for binary traits. We performed the cross-validation for all independent traits in those methods: clumping and P value thresholding (C+PT) implemented in PLINK 1.9 software, SBayesR^69^, SBayesRC, LDpred2^43^ and LDpred-funct^31^. SBayesR and LDpred2 were only run in 1M HapMap3 common SNPs due to high computational burden. C+PT, LDpred-funct and SBayesRC were run in both 1M and 7M common SNP sets. The functional annotation for LDpred-funct and SBayesR-AL was from BaseLine model 2.2 (from LDpred-funct) which contains 96 annotations, of which 14 are MAF related (10 MAFbins, MAF_Adj_Predicted_Allele_Age, MAF_Adj_ASMC and MAF_Adj_LLD_AFR). We calculated the PGS using genotypes from independent validation set in each fold and obtained the prediction R^2^ from linear regression of true phenotype on the PGS from each method for quantitative trait and McFadden’s pseudo-R^2^ for logistic regression. The final R^2^ was subtracted the value from full model (PGS + sex + age + 10 PC) to null model (sex + age + 10PC). Then, we calculated the relative prediction accuracy by 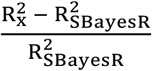, where x is each method compared and R^2^ is the prediction accuracy, then the relative prediction accuracy values are averaged across ten folds.

### Trans-ancestry prediction

We used the summary statistics from all European unrelated samples as the training data (sample sizes showed in **Supplement Table 1**). The prediction accuracy was calculated as in the within European ancestry prediction by comparing the null model without PGS and the full model with PGS. We randomly selected 500 tuning samples for methods requiring a tuning step and excluded them from the PGS calculation in all methods. The settings for different methods used in the study are described as below.

#### SBayesR family

We ran SBayesR and SBayesRC using the summary statistics of UKB EUR, and then applied the estimated SNP effects directly to the genotypes of individual from SAS, EAS and AFR ancestries in the UKB.

#### PolyPred-S

We followed the steps in ref^15^ to first apply PolyFun + SuSiE in fine-mapping of 7M SNPs with functional annotations from BaseLineLF and LD data from the PolyPred program which was calculated from the UKB (a step called PolyFun-pred). Then, the mix SNP weights were optimised in 500 tuning samples of the target ancestry using both the predictors from Polyfun-pred and predictors from SBayesR (using training samples of European ancestry). We obtained the PGS from the optimised SNP weights and genotypes for each target ancestry (SAS, EAS and AFR) in the UKB.

#### PRS-CSx and SBayesRC-multi

We evaluated these two methods based on their performance in predicting individuals of EAS ancestry, because of availability of GWAS summary statistics from BioBank Japan (BBJ)^51^ in the public domain. There were 11 traits in common between our selected traits and BBJ, of which 3 traits were diseases. PRS-CSx requires 500 cases in the tunning samples but there were no sufficient EAS case/control samples for the disease traits in UKB, hence the analysis was performed in the 8 continuous traits (**Supplementary Table 5**). The 500 hold-out samples from UKB EAS were used as the tunning sample. We matched the SNPs in the summary data from BBJ and UKB and removed SNPs with MAF < 0.005 in either population. After QC, 4,906,538 SNPs remained with functional annotations, of which 1,011,961 SNPs were in the HapMap3 panel. We ran PRS-CSx with GWAS summary statistics from UKB EUR and BBJ populations and LD data from UKB EUR and EAS downloaded from the website (https://github.com/getian107/PRScsx), using the default parameters. After obtaining the SNP weights specific to EUR and EAS populations, we calculated the EUR-based and EAS-based PGS with genotypes from UKB EAS, and then estimated the optimal weights to combine the two sets of PGS using the EAS tuning samples with phenotypes, as suggested by PRS-CSx.

The prediction accuracy was calculated from the optimised PGS and phenotypes in the remaining EAS validation samples. Followed the same strategy from PRS-CSx, we extended our method to utilise GWAS data from multiple populations (SBayesRC-multi). Instead of joint modelling, we run SBayesRC with summary statistics from UKB EUR and LD from UKB EUR, and run SBayesRC with summary statistics from BBJ and LD from UKB EAS separately. Then, the final SNP effects for EAS individuals were derived by combining the EUR- and EAS-based PGS in the EAS tuning samples as described above.

### Simulations

We performed two sets of simulations, one in 1M HapMap3 SNPs, and another in 7M imputed SNPs, following the BayesR model which allows the SNPs effects have different variance. We randomly selected 10,000 variants from the whole genome as causal variants, of which 6,000 had small effects sampled from N(0, 0.01), 30 had medium effects sampled from N(0, 0.1), and 10 variants had large effects following N(0, 1). Then, the effect sizes were scaled to give a trait heritability of 0.5. We repeated the simulation 10 times, with 10 sets of different causal variants. We also simulated a case with a trait heritability of 0.1 and the same number of causals. For the simulation scenario incorporating annotation data, we used the annotation effects estimated from height in real data analysis and calculated the per-SNP probability of membership in each mixture component by probit link function, then sampled the causal effects from that distribution.

## Data Availability

The UK Biobank data are available through formal application to the UK Biobank (http://www.ukbiobank.ac.uk). All the other datasets used in this study are available in the public domain.

## Code Availability

SBayesRC is implemented in a publicly available software GCTB at https://cnsgenomics.com/software/gctb/#Download and a R package at https://github.com/zhilizheng/SBayesRC.

## Acknowledgements

We thank D. Stetner and the Delta cluster maintenance team from the Institute for Molecular Bioscience in the University of Queensland for their support in this research. We thank Y. Wang for assistance in obtaining the ancestry information from UK Biobank. We thank J. Revez, V. Hivert and H. Wang for assistance in processing the phenotypes. Z. Zheng thanks Y. Wu and X. Hu for helpful discussions. This research was supported by the Australian National Health and Medical Research Council (1177268). This study makes use of data from the UK Biobank (project ID: 12505) and the Lifelines Biobank. A full list of acknowledgements to these data sets can be found in the **Supplementary Note**.

## Author Contributions

J.Z. conceived and supervised the study. J.Z., Z.Z. and M.E.G. developed the methods and algorithms. J.Z., Z.Z. and P.M.V. designed the experiment. Z.Z. conducted all analyses with the assistance or guidance from S.L., J.S., L.Y., P.T., J.Y., N.R.W., M.E.G., P.M.V. and J.Z. Z.Z., S.L. and J.Z. developed the software and R package. A.A., R.W., I.N. and H.S. provided the Lifelines Cohort Study data. Z.Z. and J.Z. wrote the manuscript with the participation of all authors. All the authors approved the final version of the manuscript.

