## Supplementary Note and Figures for "Leveraging functional genomic annotations and genome coverage to improve polygenic prediction of complex traits within and between ancestries"

#### Ethics approval

The University of Queensland Human Research Ethics Committee B (2011001173) approved the study.

#### Summary-data-based low-rank model

We derived a summary-data-based low-rank model from a general form of individual-level linear regression. Consider model

$$\mathbf{y} = \mathbf{X}\boldsymbol{\beta} + \mathbf{e} \quad (1)$$

where $\mathbf{y}$ is the vector of trait phenotypes adjusted for covariates, such as sex, age and principal components (PCs), $\mathbf{X}$ is the genotype matrix of $m$ SNPs standardised to have column mean zero and variance one, $\boldsymbol{\beta}$ is the vector of true SNP effects, and $\mathbf{e}$ are the residuals with $Var(\mathbf{e}) = \mathbf{I}\sigma_e^2$. Let $N$ be the sample size. Multiplying both sides of the equation by $\frac{1}{N}\mathbf{X}'$ gives

$$\frac{1}{N}\mathbf{X}'\mathbf{y} = \frac{1}{N}\mathbf{X}'\mathbf{X}\boldsymbol{\beta} + \frac{1}{N}\mathbf{X}'\mathbf{e}$$

The left-hand side is the GWAS marginal effect estimates $\mathbf{b}$. Let $\mathbf{R} = \frac{1}{N}\mathbf{X}'\mathbf{X}$ be the LD correlation matrix. Then, we have

$$\mathbf{b} = \mathbf{R}\boldsymbol{\beta} + \frac{1}{N}\mathbf{X}'\mathbf{e}$$

This is the summary-data-based model underlying many methods. Of note, in this model, the residuals have a variance-covariance structure proportional to the LD matrix, i.e., $Var\left(\frac{1}{N}\mathbf{X}'\mathbf{e}\right) = \frac{1}{N}\mathbf{R}\sigma_e^2$. It is often neither feasible nor necessary to compute the whole-genome LD matrix in humans. Alternatively, we compute $\mathbf{R}$ for each of the LD blocks that are found to be approximately independent in the human population. In this case, the genome-wide LD matrix is a block-diagonal matrix with blocks defined by LD blocks. The eigen-decomposition of $\mathbf{R}_i$ for block $i$, which can be performed independently and in parallel between block, is (the subscript is ignored for simplicity in notation)

$$\mathbf{R} = \mathbf{U}\boldsymbol{\Lambda}\mathbf{U}'$$

where $\mathbf{U}$ is the matrix of eigenvectors and $\boldsymbol{\Lambda}$ is the diagonal matrix of eigenvalues. Substitution of $\mathbf{R}$ in the equation above gives

$$\mathbf{b} = \mathbf{U}\boldsymbol{\Lambda}\mathbf{U}'\boldsymbol{\beta} + \frac{1}{N}\mathbf{X}'\mathbf{e}$$

Multiplying both sides by $\mathbf{\Lambda}^{-\frac{1}{2}}\mathbf{U}'$ gives

$$\mathbf{\Lambda}^{-\frac{1}{2}}\mathbf{U}'\mathbf{b} = \mathbf{\Lambda}^{\frac{1}{2}}\mathbf{U}'\boldsymbol{\beta} + \frac{1}{N}\mathbf{\Lambda}^{-\frac{1}{2}}\mathbf{U}'\mathbf{X}'\mathbf{e}$$

or simply,

$$\mathbf{w} = \mathbf{Q}\boldsymbol{\beta} + \boldsymbol{\epsilon} \tag{2}$$

where $\mathbf{w} = \mathbf{\Lambda}^{-\frac{1}{2}}\mathbf{U}'\mathbf{b}$ is a linear combination of marginal SNP effect estimates, $\mathbf{Q} = \mathbf{\Lambda}^{\frac{1}{2}}\mathbf{U}'$ is the new coefficient matrix, and the new residuals $\boldsymbol{\epsilon} = \frac{1}{N}\mathbf{\Lambda}^{-\frac{1}{2}}\mathbf{U}'\mathbf{X}'\mathbf{e}$ are independently and identically distributed, i.e.,

$$Var(\boldsymbol{\epsilon}) = \frac{1}{N}\mathbf{\Lambda}^{-\frac{1}{2}}\mathbf{U}'\mathbf{X}'\mathbf{X}\mathbf{U}\mathbf{\Lambda}^{-\frac{1}{2}}\frac{1}{N}$$

$$= \frac{1}{N}\mathbf{I}\sigma_\epsilon^2$$

Due to LD between SNPs and limited sample size, the LD matrix estimated from a reference sample is often rank deficient. In this case, a number of eigenvalues are zero. Additionally, small eigenvalues are subject to sampling variation in LD between GWAS and LD reference samples. To this end, we partition $\mathbf{\Lambda}$ into

$$\mathbf{\Lambda} = \begin{bmatrix} \mathbf{\Lambda}_q & \mathbf{0} \\ \mathbf{0} & \mathbf{\Lambda}_0 \end{bmatrix}$$

where $\mathbf{\Lambda}_q$ contains $q$ eigenvalues in descending order that cumulatively explain at least a given proportion ($\rho$) of variance in LD, i.e., $\rho = \frac{\sum_{i=1}^{q}\Lambda_i}{\sum_{i=1}^{m}\Lambda_i}$ where $\Lambda_i$ is the $i^{\text{th}}$ nonzero eigenvalue, and $\mathbf{\Lambda}_0$ contains remaining eigenvalues including zeros. Then the model can be written as

$$\begin{bmatrix} \mathbf{w}_q \\ \mathbf{w}_0 \end{bmatrix} = \begin{bmatrix} \mathbf{Q}_q \\ \mathbf{Q}_0 \end{bmatrix}\boldsymbol{\beta} + \begin{bmatrix} \boldsymbol{\epsilon}_q \\ \boldsymbol{\epsilon}_0 \end{bmatrix}$$

To remove the noise in LD, we discard the equations for $\mathbf{w}_0$, resulting in a low-rank model:

$$\mathbf{w}_q = \mathbf{Q}_q\boldsymbol{\beta} + \boldsymbol{\epsilon}_q$$

where $\mathbf{Q}_q$ has a dimension of $q \times m$ with $q \ll m$. In essence, the true SNP effects are fitted to a smaller number of effective data points rather than the observed data points that are highly correlated to each other. This model is general and can be applied with different assumptions on the distribution of SNP effects $\boldsymbol{\beta}$.

**Alternative parameterization for $\boldsymbol{\pi}_j$**

To remove the dependence between elements of $\boldsymbol{\pi}_j$ for each SNP, we employed an alternative parameterization for modelling membership probabilities and annotation effects. Let $\delta_j$ be the indicator for the mixture component membership for SNP $j$:

$$\delta_j = k \text{ with probability } \pi_{jk}; k = 1 \text{ to } 5$$

We define a conditional probability that the SNP effect belongs to the $k^{\text{th}}$ distribution given that it has passed the bar for the $(k\text{-}1)^{\text{th}}$ distribution as

$$p_{jk} = \Pr(\delta_j \geq k \mid \delta_j \geq k-1) \text{ for } k \geq 2$$

such that

$$\begin{aligned}
\pi_{j1} &= 1 - p_{j2} \\
\pi_{j2} &= (1 - p_{j3})p_{j2} \\
\pi_{j3} &= (1 - p_{j4})p_{j3}p_{j2} \\
\pi_{j4} &= (1 - p_{j5})p_{j4}p_{j3}p_{j2} \\
\pi_{j5} &= p_{j5}p_{j4}p_{j3}p_{j2}
\end{aligned}$$

We then apply the generalised linear model to link $p_{jk}$ with $\boldsymbol{\alpha}_k$, i.e.,

$$g(p_{jk}) = \mu_k + \sum_{c=1}^{C} A_{jc}\alpha_{kc}$$

In this parameterisation, all $p_{jk}$ are independent, which means that $\boldsymbol{\alpha}_k$ can be sampled in parallel in each MCMC iteration, and $\alpha_{kc}$ can be sampled from its full conditional distribution using Gibbs sampling algorithm, following the algorithm of Albert and Chib[1].

Let $z_{jk}$ be the indicator variable for whether the SNP effect can "climb" up to a higher distribution, i.e.,

$$z_{jk} \sim Bernoulli(p_{jk})$$

To allow Gibbs sampling, a probit link function is chosen, namely $g^{-1}(p_{jk}) = \Phi(p_{jk})$ where $\Phi(\cdot)$ is the cumulative density function (CDF) of the standard normal distribution. It has been shown that with an auxiliary variable $l_{jk}$ defined as

$$z_{jk} = \begin{cases} 0, & l_{jk} > 0 \\ 1, & l_{jk} \leq 0 \end{cases}$$

a linear model can be constructed

$$l_{jk} = \mu_k + \sum_{c=1}^{C} A_{jc}\alpha_{kc} + \varepsilon_{jk}$$

with $\varepsilon_{jk} \sim N(0,1)$. In this model, given a normal prior distribution, $\alpha_{kc} \sim N(0, \sigma^2_{\alpha_k})$, the full conditional distribution for $\alpha_{kc}$ is a univariate normal distribution, since $\alpha_{kc}$ is conditionally independent of $\boldsymbol{z}_k$ given $\boldsymbol{l}_k$. Given $z_{jk}$ and $\boldsymbol{\alpha}_k$, the full conditional distribution for the latent variable $l_{jk}$ is a truncated normal distribution. The Gibbs sampling procedure is described in the following section.

**Scaling the SNP marginal effect estimates**

The derivation for the summary-data-based model is based on the marginal effects in units of per standardized genotype (**b**). When the marginal effects were estimated from GWAS using genotypes at 0/1/2 scale (**b***), **b** can be estimated, in a scalar form, by

$$b_j = s_j b_j^* \text{ where } s_j = \sqrt{\frac{\sigma_y^2}{N_j\sigma_j^2 + \left(b_j^*\right)^2}}$$

where $\sigma_y^2$ is the phenotypic variance, $N_j$ is the per-SNP sample size, and $\sigma_j$ is the standard error for SNP $j$. If the trait phenotypes are not standardized, the phenotypic variance can be estimated by taking the median value of $2f_j(1-f_j)\left[N_j\sigma_j^2 + \left(b_j^*\right)^2\right]$ across SNPs, where $f_j$ is the allele frequency in the GWAS sample (ref[2,3]). The per-SNP sample size $N_j$ can be replaced by the overall sample size *N*. Here, we assume $\sigma_y^2 = 1$, then

$$s_j = \sqrt{\frac{1}{N_j\sigma_j^2 + \left(b_j^*\right)^2}}$$

and will scale the joint effect estimate $\beta_j$ back to the phenotypic scale using the same $s_j$ so that this parsimonious assumption would not have an impact on the result (ref.[4,5]).

**Violation of model assumptions**

There are at least two important assumptions implied in the general form of summary-data-based models[5]. One assumption is that the LD correlation matrix calculated from the reference sample is consistent with that from the GWAS sample, which is violated when the LD reference has a too small sample size (i.e., large sampling variation in LD) or is genetically different from the GWAS sample. Another assumption is that the summary statistics are derived from the same set of individuals for all SNPs, which may not hold when the summary statistics are obtained from a meta-analyses where different SNP genotyping panels, imputation references or quality control (QC) procedures are used in different cohorts. Failure to satisfy these assumptions can result in severe model misspecifications. In

SBayesRC (or SBayesRC without annotations), we aim to account for the heterogeneity in both LD and per-SNP sample size by removing those principal components with the smallest eigenvalues in the LD matrix and estimating the residual variance from the data (**Methods**). We performed genome-wide simulations based on the imputed SNP data in the UKB to assess the robustness of our method to model misspecifications, in comparison of state-of-the-art methods including LDpred2[4] and SBayesR.

**Estimation of residual variance helps to improve model robustness**

In the summary-data-based model, Eq (1), $Var(\boldsymbol{\varepsilon}) = \frac{1}{N}\mathbf{R}\sigma_\epsilon^2$. It is often assumed that $\sigma_\epsilon^2 \approx \sigma_y^2$ given a negligible proportion of variance explained by a single SNP, and further $\sigma_\epsilon^2 \approx 1$ assuming a unit phenotypic variance for the trait. It is possible, however, that $\sigma_\epsilon^2 > 1$ if there exist large LD differences between GWAS and LD reference samples. This is because using summary statistics from GWAS and inaccurate LD data from a reference is analogous to using estimated genotype data with noise in the fitted model for GWAS:

$$
\begin{aligned}
\mathbf{y} &= \mathbf{1}\mu + \hat{\mathbf{X}}\hat{b} + \mathbf{e} \\
&= \mathbf{1}\mu + (\mathbf{X} + \boldsymbol{\Delta})\hat{b} + \mathbf{e} \\
&= \mathbf{1}\mu + \mathbf{X}\hat{b} + \left(\boldsymbol{\Delta}\hat{b} + \mathbf{e}\right) \\
&= \mathbf{1}\mu + \mathbf{X}\hat{b} + \mathbf{e}^*
\end{aligned}
$$

where $\hat{\mathbf{X}}$ is the combination of genotypes used in GWAS (**X**) and the differences to those observed from the reference ($\boldsymbol{\Delta}$), and $\hat{b}$ is the ordinary least squares estimate for the SNP effect. It can be seen that the new residual in the above model can have variance larger than the phenotypic variance when the noise in the genotype data is large, i.e., $Var(\Delta\hat{b} + \mathrm{e}) > \mathrm{Var}(y)$. Thus, it would be beneficial to estimate the residual variance from the data given the GWAS summary statistics and reference LD data.

In contrast to Eq (1), it is very straightforward to estimate the residual variance in the low-rank model, Eq (2), because the residuals are independently distributed, $Var(\boldsymbol{\epsilon}) = \frac{1}{N}\mathbf{I}\sigma_\epsilon^2$. The MCMC sampling process for the residual variance is shown as below. The large residual variance estimate will introduce a shrinkage mechanism to manage the potential convergence issue due to violation of model assumptions. It has been found that SNP effect sizes would blow up during MCMC when the model fails to converge. In this case, the sampled values of residual variances would be large if the SNP effect sizes tend to blow up, which will in turn shrink them back toward zero, preventing the failure in convergence.

**Estimation of SNP-based heritability and per-SNP heritability enrichment for each annotation**

The total genetic variance is

$$\begin{aligned}\sigma_g^2 &= \boldsymbol{\beta}'\mathbf{R}\boldsymbol{\beta} \\ &= \boldsymbol{\beta}'\mathbf{U}\boldsymbol{\Lambda}\mathbf{U}'\boldsymbol{\beta} \\ &= \boldsymbol{\beta}'\mathbf{U}\boldsymbol{\Lambda}^{\frac{1}{2}}\boldsymbol{\Lambda}^{\frac{1}{2}}\mathbf{U}'\boldsymbol{\beta} \\ &= \boldsymbol{\beta}'\mathbf{Q}\mathbf{Q}\boldsymbol{\beta} \\ &= \hat{\mathbf{w}}'\hat{\mathbf{w}}\end{aligned}$$

We calculate this quantity in each of MCMC iterations given the sampled values of SNP effects $\boldsymbol{\beta}$. Assuming unit phenotypic variance, the SNP-based heritability $h_{\mathrm{SNP}}^2 = \sigma_g^2$, estimated by the posterior mean of MCMC samples discarding the samples from the burn-in period.

For a binary annotation $c$, the total variance explained by the SNPs within the annotation is calculated as

$$\sigma_c^2 = \sum_{j=1}^{m_c} \beta_{jc}^2$$

where $m_c$ is the number of SNPs within the annotation. The per-SNP heritability enrichment ($\theta_c$) is then calculated as

$$\theta_c = \frac{\sigma_c^2}{m_c} \Big/ \frac{\sigma_g^2}{m}$$

For a quantitative annotation, the per-SNP heritability enrichment is calculated as the slope of the regression of $\beta_{jc}^2$ on the annotation value $A_{jc}$

$$\mathrm{E}[\boldsymbol{\beta}_c^2] = \mathbf{1}\mu_c + \mathbf{A_c}\omega_c$$

and $\theta_c = 1 + \omega_c$. Similarly, we compute $\theta_c$ in every iteration of MCMC and estimate by the posterior mean after burn-in.

**MCMC sampling scheme**

We use MCMC sampling to draw posterior inference on the model parameters. The joint distribution of data and all parameters in the low-rank model is

$$f(\mathbf{w},\boldsymbol{\beta},\boldsymbol{\pi},\mathbf{z},\boldsymbol{\alpha},\boldsymbol{\sigma}_{\alpha}^{2},\sigma_{\epsilon}^{2}) \propto (\sigma_{\epsilon}^{2})^{-\frac{q}{2}} \exp\left\{-\frac{(\mathbf{w}-\mathbf{Q}\boldsymbol{\beta})'(\mathbf{w}-\mathbf{Q}\boldsymbol{\beta})}{2\frac{\sigma_{\epsilon}^{2}}{n}}\right\}$$

$$\times \prod_{j=1}^{m}\left\{\sum_{k=1}^{5}\pi_{jk}\left[\exp\left\{-\frac{\beta_j^2}{2\gamma_k\sigma_g^2}\right\}\right]\right\}$$

$$\times \prod_{k=2}^{5}\prod_{j=1}^{m}\Phi\left(\mu_k+\mathbf{A}_j'\boldsymbol{\alpha}_k\right)^{z_{jk}}\left[1-\Phi\left(\mu_k+\mathbf{A}_j'\boldsymbol{\alpha}_k\right)\right]^{(1-z_{jk})}$$

$$\times \prod_{k=2}^{5}\prod_{c=1}^{C}\left(\sigma_{\alpha_c}^2\right)^{-\frac{1}{2}}\exp\left\{-\frac{\alpha_{kc}^2}{2\sigma_{\alpha_c}^2}\right\}$$

$$\times \prod_{k=2}^{5}\left(\sigma_{\alpha_c}^2\right)^{-\frac{2+v_\alpha}{2}}\exp\left\{-\frac{v_\alpha\tau_\alpha^2}{2\sigma_{\alpha_c}^2}\right\}$$

$$\times (\sigma_{\epsilon}^{2})^{-\frac{2+v_\epsilon}{2}}\exp\left\{-\frac{v_\epsilon\tau_\epsilon^2}{2\sigma_\epsilon^2}\right\}$$

Suppose $\delta_j$ is the indicator variable to the distribution membership of $\beta_j$. The full conditional distribution for $\beta_j$ is

$$f\left(\beta_j\middle|\mathbf{w},\boldsymbol{\beta}_{-j},\delta_j,\sigma_g^2,\sigma_\epsilon^2\right)$$

$$\propto (\sigma_{\epsilon}^{2})^{-\frac{q}{2}} \exp\left\{-\frac{\left(\mathbf{w}-\sum_{j'\neq j}\mathbf{Q}_{j'}\beta_{j'}\right)'\left(\mathbf{w}-\sum_{j'\neq j}\mathbf{Q}_{j'}\beta_{j'}\right)}{2\frac{\sigma_{\epsilon}^{2}}{n}}\right\}\exp\left\{-\frac{\beta_j^2}{2\gamma_k\sigma_g^2}\right\}$$

$$= N\left(\frac{r_j}{C_j},\frac{\sigma_\epsilon^2}{C_j}\right)$$

where

$$r_j = \mathbf{Q}_j'\left(\mathbf{w}-\sum_{j'\neq j}\mathbf{Q}_{j'}\beta_{j'}\right)$$

$$= \mathbf{Q}_j'\mathbf{e}+\beta_j$$

$$C_j = 1+\frac{\sigma_\epsilon^2}{\gamma_k\sigma_g^2}$$

The full conditional distribution for $\delta_j$ is

$$\Pr\left(\delta_j=k\middle|\mathbf{w},\boldsymbol{\beta},\sigma_g^2,\sigma_\epsilon^2\right)=\frac{f\left(\mathbf{w}\middle|\delta_j=k,\boldsymbol{\beta},\sigma_g^2,\sigma_\epsilon^2\right)f\left(\delta_j=k\right)}{\sum_{k'=1}^{5}f\left(\mathbf{w}\middle|\delta_j=k',\boldsymbol{\beta},\sigma_g^2,\sigma_\epsilon^2\right)f\left(\delta_j=k'\right)}$$

where $f\left(\mathbf{w}\middle|\delta_j=k,\boldsymbol{\beta},\sigma_g^2,\sigma_\epsilon^2\right)$ is shown as above and $f\left(\delta_j=k\right)=\pi_k$.

As described above, $\boldsymbol{\pi}_j$ is a function of $\boldsymbol{p}_j$, and $\boldsymbol{p}_j$ is a linear model of $\boldsymbol{\alpha}_k$ through the probit link, i.e.,

$$p_{jk} = \Phi^{-1}\left(\mu_k + \mathbf{A}_j'\boldsymbol{\alpha}_k\right)$$

Here, we introduce another indicator variable $z_{jk}$, where

$$z_{jk} = 1 \text{ if } \delta_j = k \text{ for } k \geq 2$$

and a latent variable $l_{jk}$, for which the full conditional distribution is

$$l_{jk}|z_{jk}, \mu_k, \boldsymbol{\alpha}_k = \begin{cases} TN\left(\mu_k + \mathbf{A}_j'\boldsymbol{\alpha}_k, 1, 0, \infty\right), & \text{if } z_{jk} = 1 \\ TN\left(\mu_k + \mathbf{A}_j'\boldsymbol{\alpha}_k, 1, -\infty, 0\right), & \text{if } z_{jk} = 0 \end{cases}$$

Similar to the sampling of SNP effects, we use single-site Gibbs sampler to sample the annotation effects. The full conditional distribution for $\alpha_{kc}$ is

$$\begin{aligned} &f\left(\alpha_{kc}|\boldsymbol{l}_k, \boldsymbol{\alpha}_{-kc}, \sigma_{\alpha_k}^2\right) \\ &\quad \propto \exp\left\{-\frac{1}{2}\left(\boldsymbol{l}_k - \sum\nolimits_{c' \neq c} \mathbf{A}_{c'}\alpha_{kc'}\right)'\left(\boldsymbol{l}_k - \sum\nolimits_{c' \neq c} \mathbf{A}_{c'}\alpha_{kc'}\right)\right\} \exp\left\{-\frac{\alpha_{kc}^2}{2\sigma_{\alpha_k}^2}\right\} \\ &\quad = N\left(\frac{r_{kc}}{C_{kc}}, \frac{1}{C_{kc}}\right) \end{aligned}$$

where

$$r_{kc} = \mathbf{A}_c'\left(\boldsymbol{l}_k - \sum\nolimits_{j' \neq j} \mathbf{A}_{c'}\alpha_{kc'}\right)$$

$$C_{kc} = \mathbf{A}_c'\mathbf{A}_c + \frac{1}{\sigma_{\alpha_k}^2}$$

The full conditional distribution for $\sigma_{\alpha_k}^2$ is

$$\begin{aligned} f\left(\sigma_{\alpha_k}^2|\boldsymbol{\alpha}_k\right) &\propto f\left(\boldsymbol{\alpha}_k|\sigma_{\alpha_k}^2\right)f\left(\sigma_{\alpha_k}^2\right) \\ &\propto \left(\sigma_{\alpha_k}^2\right)^{-\frac{C+v_\alpha+2}{2}} \exp\left\{-\frac{\boldsymbol{\alpha}_k'\boldsymbol{\alpha}_k + v_\alpha\tau_\alpha^2}{2\sigma_{\alpha_k}^2}\right\} \\ &= \chi^{-2}\left(\tilde{v}_\alpha, \tilde{\tau}_\alpha^2\right) \end{aligned}$$

where $\tilde{v}_\alpha = C + v_\alpha$ and $\tilde{\tau}_\alpha^2 = (\boldsymbol{\alpha}_k'\boldsymbol{\alpha}_k + v_\alpha\tau_\alpha^2)/\tilde{v}_\alpha$.

The full conditional distribution for $\sigma_\epsilon^2$ is

$$\begin{aligned} &f(\sigma_\epsilon^2|\mathbf{w}, \boldsymbol{\beta}) \propto f(\mathbf{w}|\boldsymbol{\beta}, \sigma_\epsilon^2)f(\sigma_\epsilon^2) \\ &\propto (\sigma_\epsilon^2)^{-\frac{q}{2}} \exp\left\{-\frac{\left(\mathbf{w} - \sum_j \mathbf{Q}_j\beta_j\right)'\left(\mathbf{w} - \sum_j \mathbf{Q}_j\beta_j\right)}{2\frac{\sigma_\epsilon^2}{n}}\right\} (\sigma_\epsilon^2)^{-\frac{v_\epsilon+2}{2}} \exp\left\{-\frac{v_\epsilon\tau_\epsilon^2}{2\sigma_\epsilon^2}\right\} \end{aligned}$$

$$\propto (\sigma_\epsilon^2)^{-\frac{q+v_\epsilon+2}{2}} \exp\left\{-\frac{\boldsymbol{\epsilon}'\boldsymbol{\epsilon} + v_\epsilon\tau_\epsilon^2}{2\sigma_\epsilon^2}\right\}$$

$$= \chi^{-2}(\tilde{v}_\epsilon, \tilde{\tau}_\epsilon^2)$$

where $\tilde{v}_\epsilon = q + v_\epsilon$ and $\tilde{\tau}_\epsilon^2 = (\boldsymbol{\epsilon}'\boldsymbol{\epsilon} + v_\epsilon\tau_\epsilon^2)/\tilde{v}_\epsilon$.

### Algorithm pseudo code

**SBayesRC algorithm**

**Input**: GWAS summary statistics, reference LD correlation matrix, functional annotation data

Scale the GWAS marginal effect estimate $b_j = s_j b_j^*$

Construct the low-rank model by performing eigen-decomposition on LD blocks

Initialize model parameters

**for** i :=1 **to** number of iterations **do**

    **for** j := 1 **to** number of SNPs **do**

        Calculate $r_j = \mathbf{Q}_j'\mathbf{w}_{corr} + \beta_j$

        Calculate $C_j = 1 + \frac{\sigma_\epsilon^2}{\gamma_k\sigma_g^2}$ for each $\gamma_k$

        Calculate the posterior probabilities of SNP effect distribution memberships and sample $\delta_j$

        Sample SNP effect $\beta_j$ from its full conditional distribution $N\left(\frac{r_j}{C_j}, \frac{\sigma_\epsilon^2}{C_j}\right)$

        Given the sampled value of $\beta_j^{new}$, adjust $\mathbf{w}_{corr}^{new} = \mathbf{w}_{corr}^{old} + \mathbf{Q}_j(\beta_j^{old} - \beta_j^{new})$

        Calculate indicator variables $\mathbf{z}_j$ given $\delta_j$

    **end**

    **for** k := 2 **to** number of mixture distribution components **do**

        **for** j := 1 **to** number of SNPs that passed the bar for current component **do**

            Sample latent variable $l_{jk}$ from a truncated normal distribution given $z_{jk}$

        **end**

        **for** c := 1 **to** number of annotations **do**

            Sample annotation effect $\alpha_{kc}$ from its full conditional distribution given $\boldsymbol{l}_k$

        **end**

        Sample annotation effect variance $\sigma_{\alpha_k}^2$ from its full conditional distribution given $\boldsymbol{\alpha}_k$

    **end**

    Calculate $\hat{\mathbf{w}} = \mathbf{Q}\boldsymbol{\beta}$ and total genetic variance $\sigma_g^2 = \hat{\mathbf{w}}'\hat{\mathbf{w}}$

    Estimate the per-SNP heritability enrichment for each annotation

    Sample residual variance $\sigma_\epsilon^2$ from its full conditional distribution for each block

**end**

Scale back the posterior mean SNP joint effects to per-allele scale by $\hat{\beta}_j^* = s_j\hat{\beta}_j$

**Summary data imputation**

We implement the imputation method for summary data from impG to avoid the heavy re-calculation of the eigen decomposition for the LD matrix if some SNPs are missing from the LD panel. The imputation is based on the Z score, correlation information among missing SNPs and the typed SNPs. The Z score for the missing SNPs can be obtained from

$$\mathbf{z}_i = \mathbf{R}_{it}\mathbf{R}_{tt}^{-1}\mathbf{Z}_\mathbf{t}$$

Where $\mathbf{z}_i$ is the imputed Z score for missing SNPs in the GWAS summary data, $\mathbf{R}_{it}$ is the LD correlation matrix among the missing SNPs and typed SNPs, $\mathbf{R}_{tt}$ is the LD correlation matrix among the typed SNPs.

We converted the $\mathbf{z}_i$ for missing SNPs to marginal effects at 0/1/2 scale ($\mathbf{b}^*$) and standard error ($\sigma_i$) by

$$\sigma_i = \frac{\sigma_y}{\sqrt{2f_i(1-f_i)(N_i + z_i^2)}}$$

$$b^* = z_i\sigma_i$$

Where $N_i$ is the per-SNP sample size (replaced by median per-SNP sample size of known SNPs instead), $f_i$ is the allele frequency from reference genotype, $\sigma_y$ is the phenotypic standard derivation (square root of phenotypic variance). The phenotypic variance can be estimated by taking the median value of $2f_i(1-f_i)[N_i\sigma_i^2 + (b_i^*)^2]$ across SNPs, here $f_i$ is the allele frequency in the GWAS sample (ref[2,3]).

**UMCG Genetics Lifelines Initiative (UGLI) group author**

LifeLines Cohort Study

Raul Aguirre-Gamboa (1), Patrick Deelen (1), Lude Franke (1), Jan A Kuivenhoven (2), Esteban A Lopera Maya (1), Ilja M Nolte (3), Serena Sanna (1), Harold Snieder (3), Morris A Swertz (1), Peter M. Visscher (3,4), Judith M Vonk (3), Cisca Wijmenga (1)

(1) Department of Genetics, University of Groningen, University Medical Center Groningen, The Netherlands

(2) Department of Pediatrics, University of Groningen, University Medical Center Groningen, The Netherlands

(3) Department of Epidemiology, University of Groningen, University Medical Center Groningen, The Netherlands

(4) Institute for Molecular Bioscience, The University of Queensland, Brisbane, Queensland, Australia.

**Acknowledgements**

Lifelines Cohort Study

The Lifelines Biobank initiative has been made possible by funding from the Dutch Ministry of Health, Welfare and Sport, the Dutch Ministry of Economic Affairs, the University Medical Center Groningen (UMCG the Netherlands), University of Groningen and the Northern Provinces of the Netherlands. The generation and management of GWAS genotype data for the Lifelines Cohort Study is supported by the UMCG Genetics Lifelines Initiative (UGLI). UGLI is partly supported by a Spinoza Grant from NWO, awarded to Cisca Wijmenga. The authors wish to acknowledge the services of the Lifelines Cohort Study, the contributing research centers delivering data to Lifelines, and all the study participants.

### Supplementary Figures

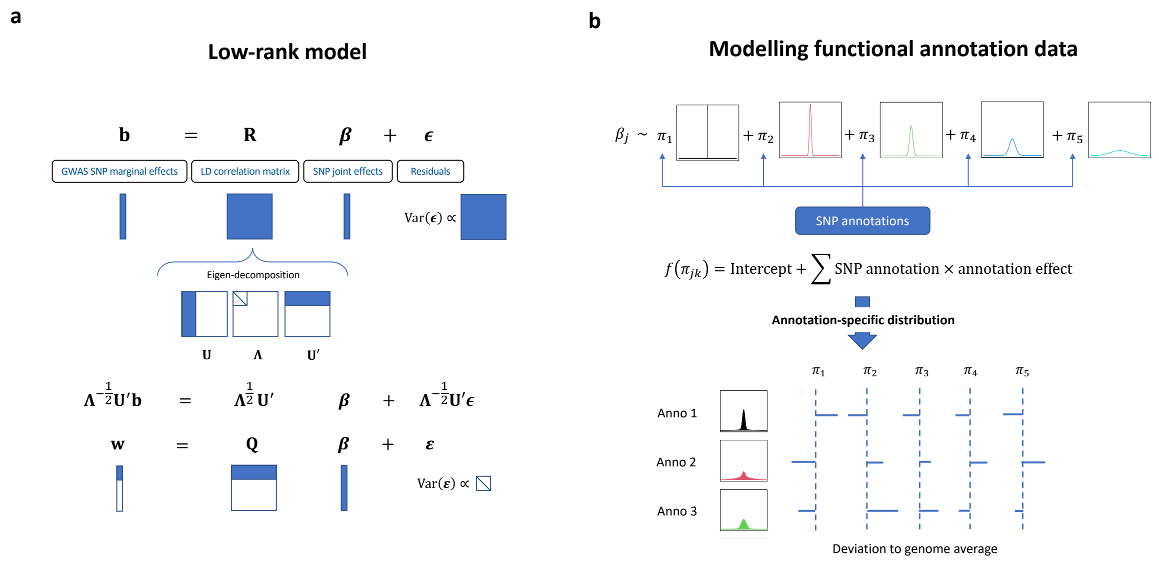


**Supplementary Figure 1** Schematic overview of SBayesRC. a) A resource-efficient low-rank model that can simultaneously fit sequence-level SNPs with high computation efficiency and has independent residuals. b) A hierarchical multi-component mixture prior for SNP effects that incorporates functional annotation data and allows for any distribution of SNP effects in each annotation.

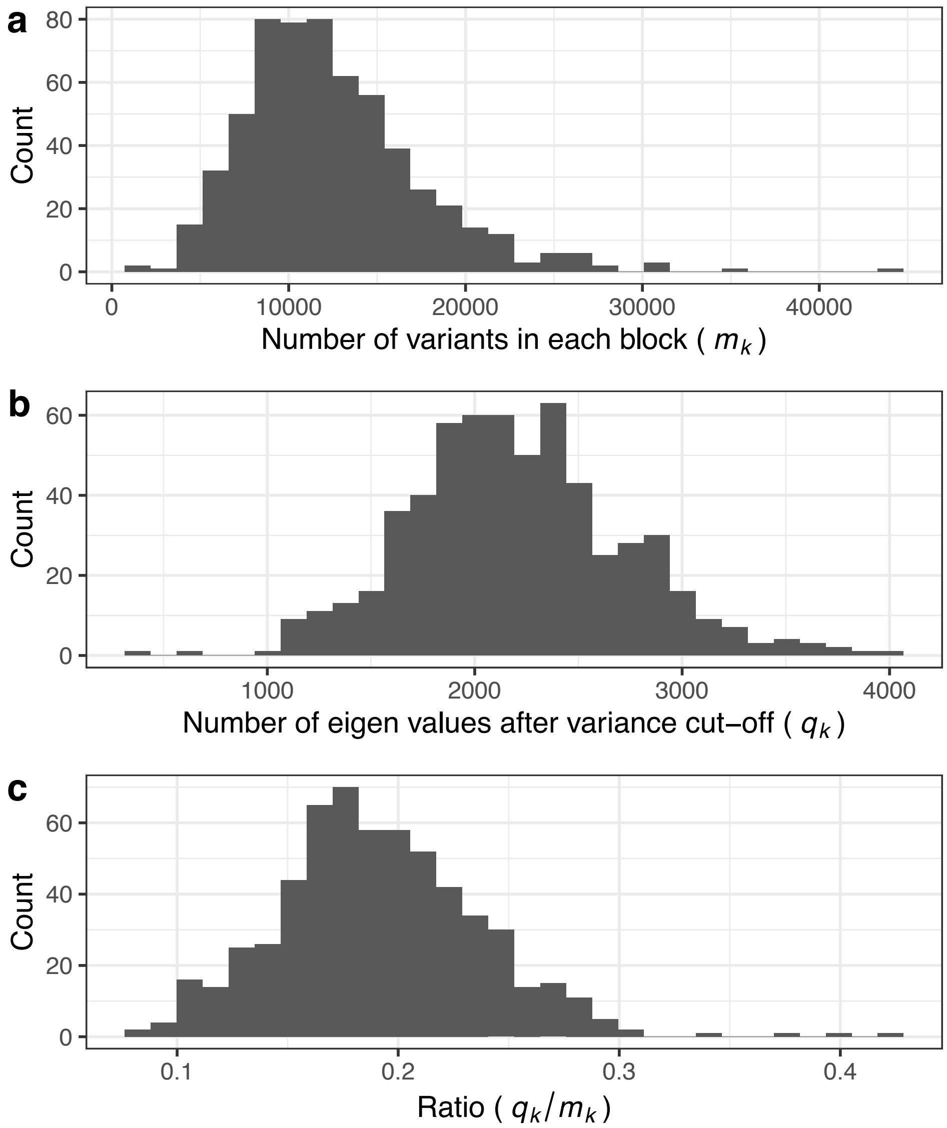


**Supplementary Figure 2** The low-rank model leads to a substantial reduction in dimension. a) The distribution of the number of SNPs per block ($m_k$). b) The distribution of the number of principal components ($q_k$) that collectively explain at least $\rho$ proportion of LD variance in each block ($\rho = 99.5\%$). c) The distribution of $q_k/m_k$ at $\rho = 99.5\%$.

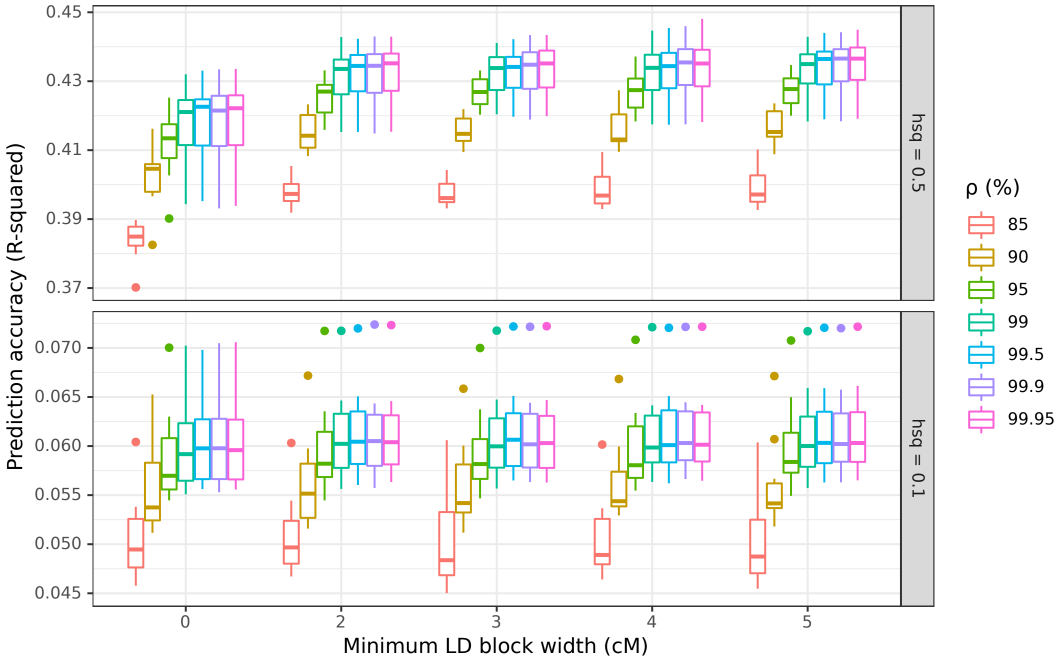


**Supplementary Figure 3** Prediction accuracy of SBayesRC using different minimum values of LD block width and different minimum proportions ($\rho$) of variance in the LD matrix in the simulated data with heritability = 0.1 or 0.5. Minimum LD block width = 0 means using the original quasi-independent LD blocks found in the European population (ref[6]) without merging of small LD blocks.

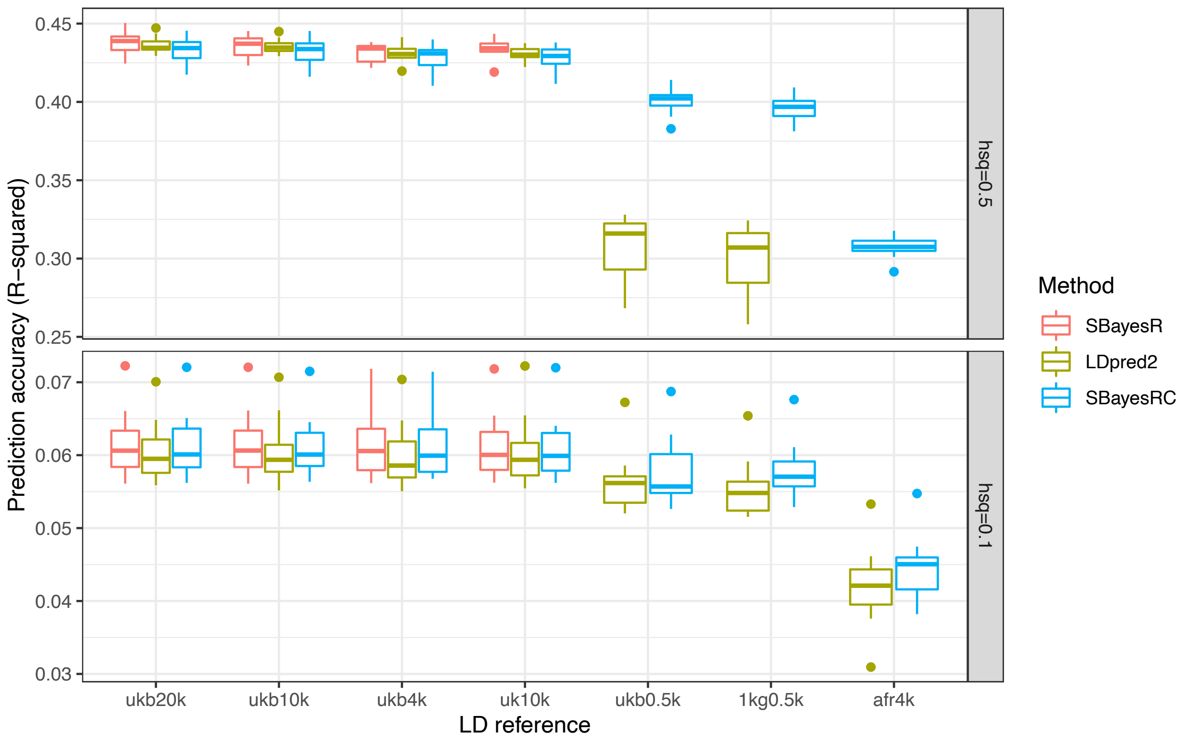


**Supplementary Figure 4** Robustness of SBayesRC to the choice of LD reference in simulation with HapMap3 SNP panel. LD reference data sets included ukb20k: 20,000 random sample from the UKB of European ancestry (EUR); ukb10k: 10,000 random sample from UKB EUR; ukb4k: 4,000 random sample from UKB EUR; uk10k: 3,642 unrelated samples from the UK10K data set; ukb0.5k: 500 random sample from UKB EUR; 1kg0.5k: 494 unrelated samples from 1000GP EUR; afr4k: 4,000 random samples from the UKB of African ancestry (AFR).

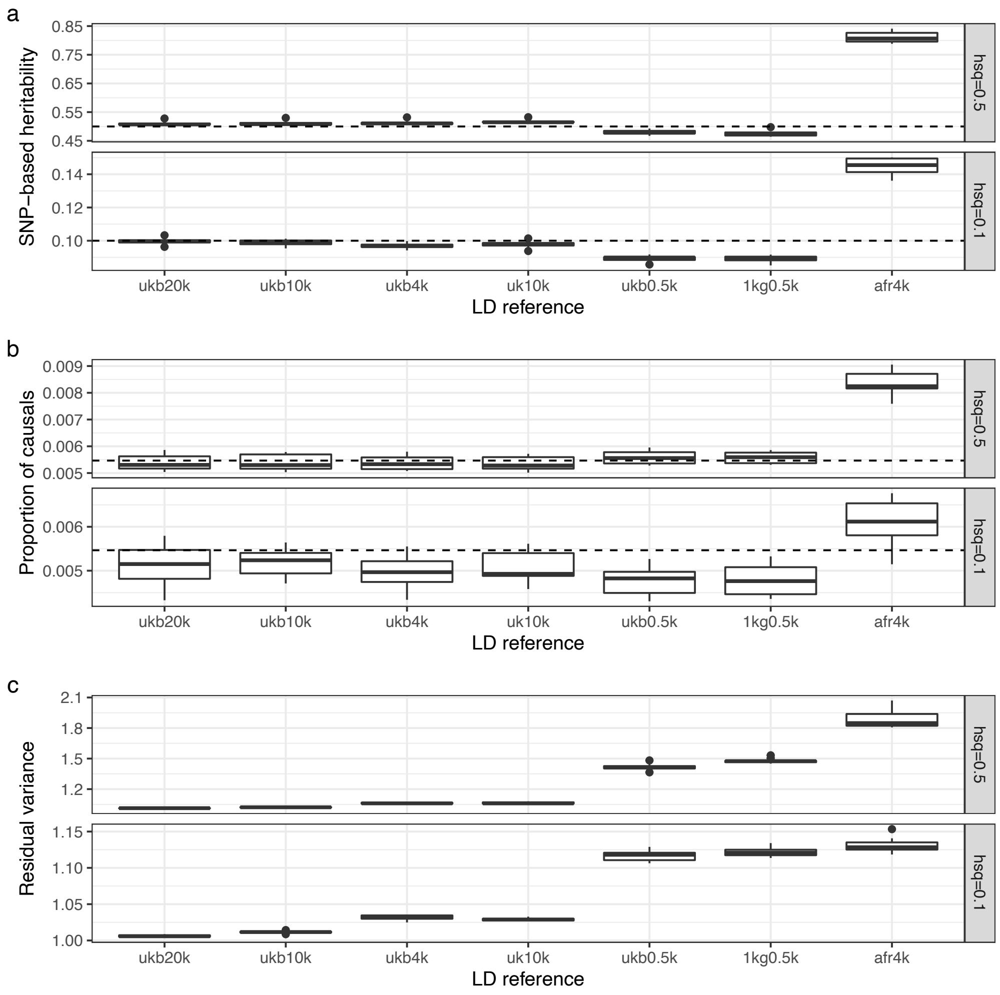


**Supplementary Figure 5** Estimation of SNP-based heritability, polygenicity (the proportion of causal variants) and residual variance in SBayesRC without annotation using1M HapMap3 SNPs and different choices of LD reference for a simulated trait with heritability = 0.1 or 0.5. The dashed line in panel a and b indicates the true value in the simulation.

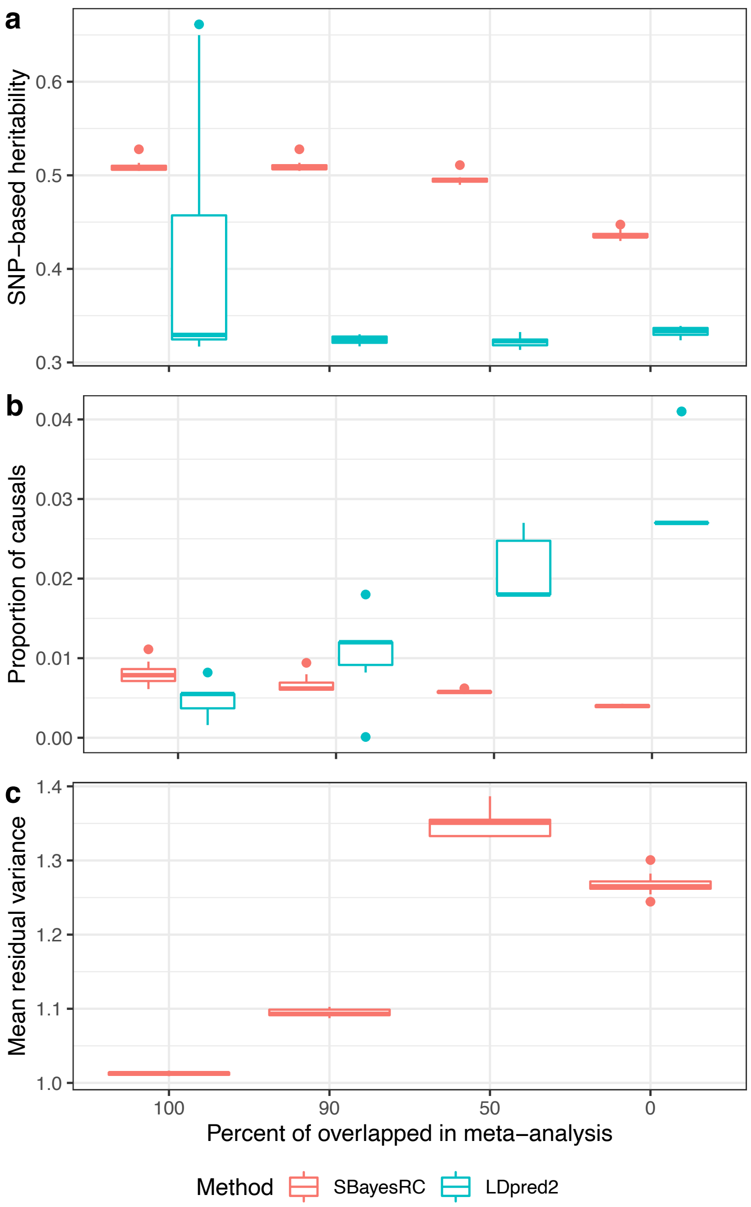


**Supplementary Figure 6** Parameter estimation from SBayesRC and LDpred2 using summary statistics from a meta-analysis of two simulated cohorts where the proportion of overlapped SNPs between the two cohorts varied from 100 to 0. The proportion of overlapping is less than 100, there existed unequal per-SNP sample sizes in the GWAS summary data. SBayesRC gave approximately unbiased estimates for SNP-based heritability (true value = 0.5) and polygenicity (true value = 0.01), whereas these estimates in LDpred2 were largely biased (panel a and b). The model misspecification affected the residual variance in SBayesRC, which is a nuisance parameter in the model (panel c).

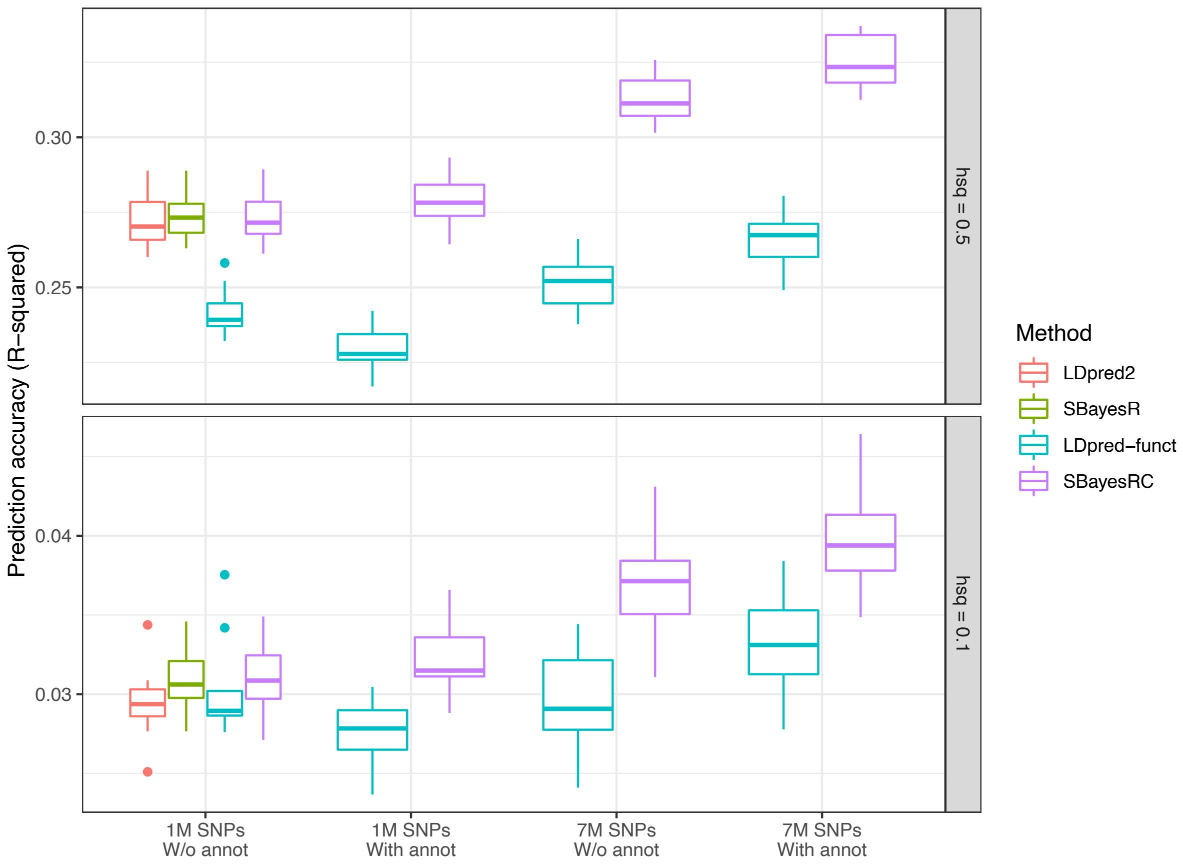


**Supplementary Figure 7** Prediction accuracy of phenotypes using PGS derived from different methods for simulated traits (heritability = 0.1 or 0.5).

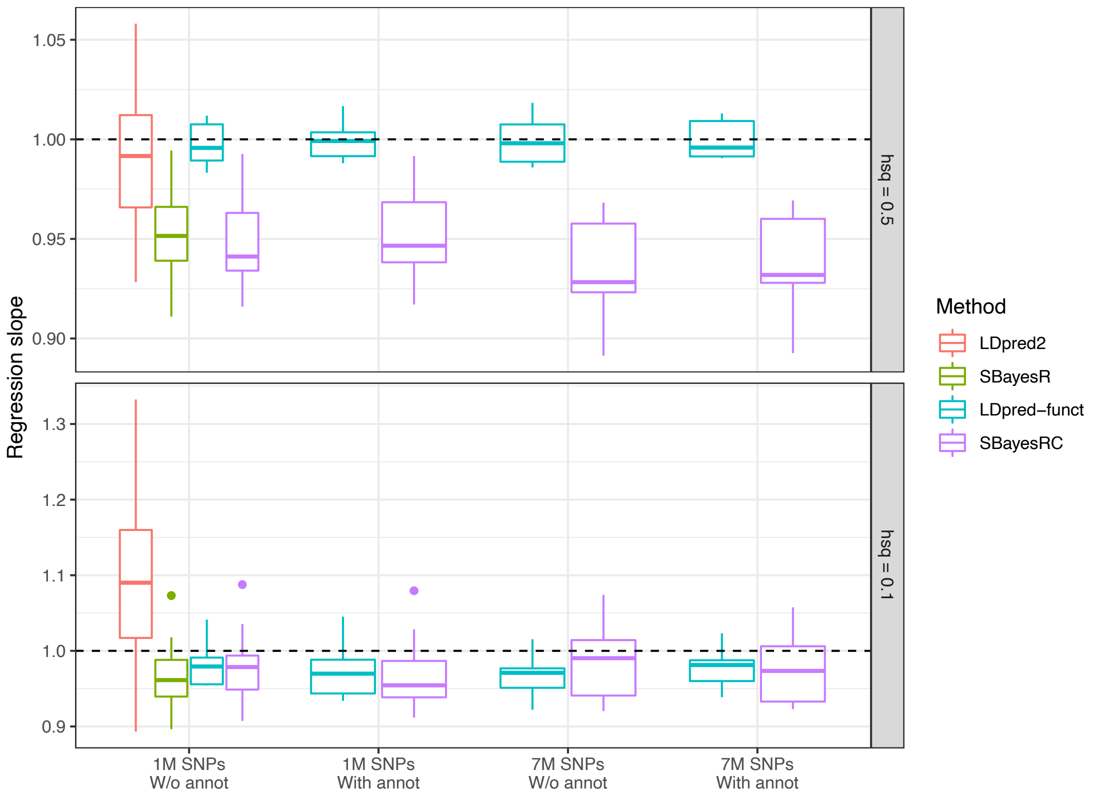


**Supplementary Figure 8** Slope of regression (bias) of phenotypes using PGS derived from different methods and SNP panels using the simulated data.

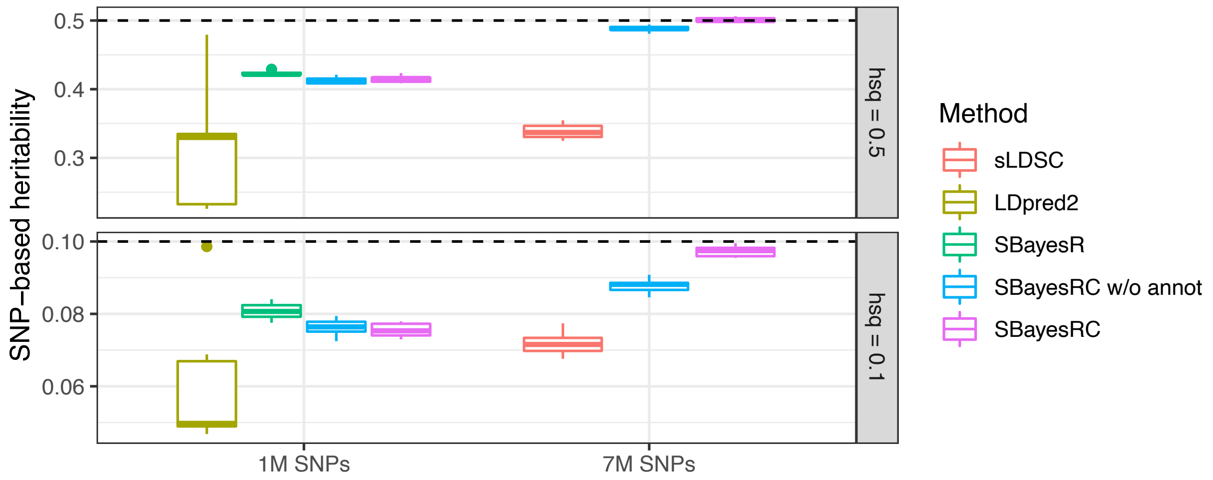


**Supplementary Figure 9** SNP-based heritability estimation from different methods and SNP panels for simulated traits with heritability = 0.1 or 0.5.

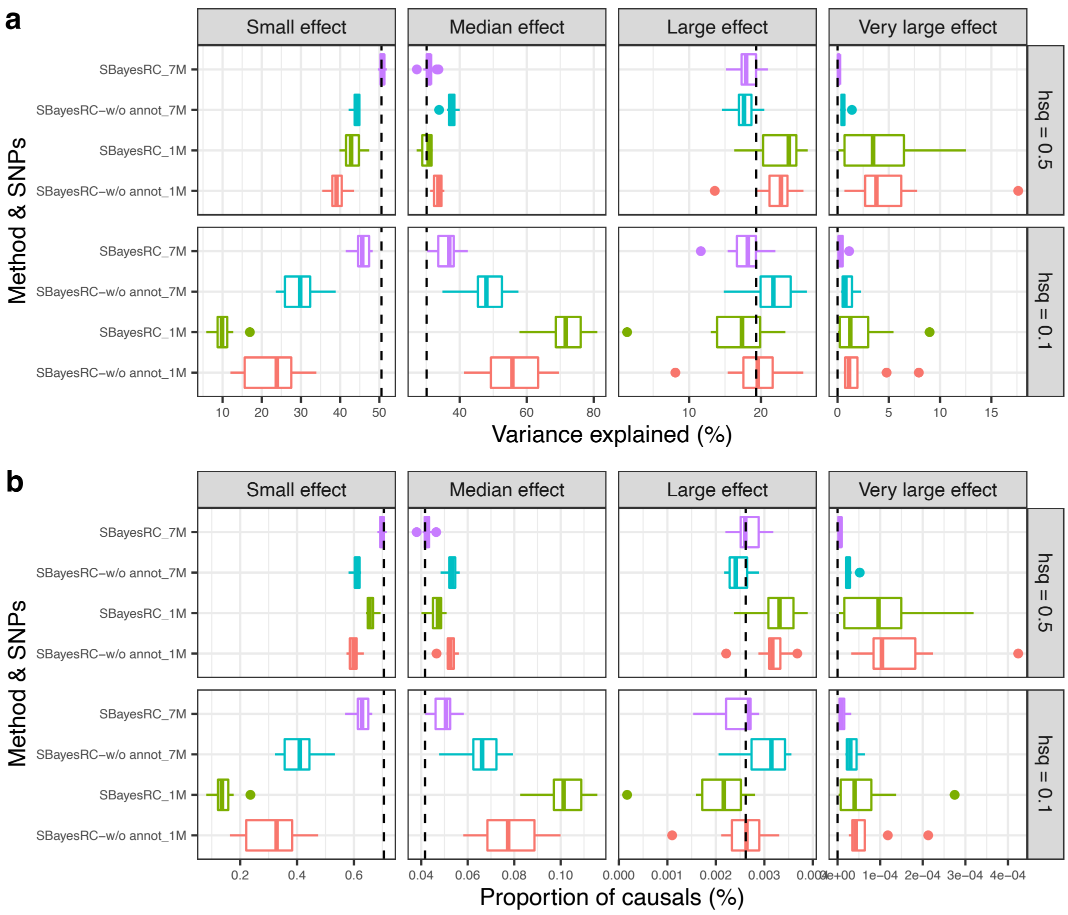


**Supplementary Figure 10** Genetic architecture parameter estimation using SBayesRC without annotation or SBayesRC (incorporating annotation data) with different SNP panels for a simulated trait (heritability = 0.1 or 0.5). The dashed line indicates the true value in the simulation.

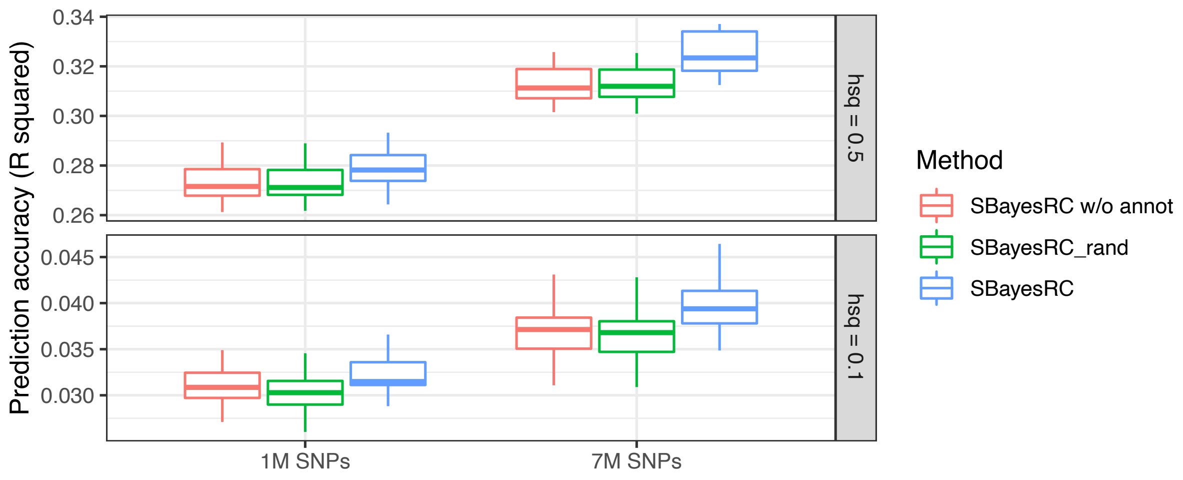


**Supplementary Figure 11** SBayesRC improved prediction accuracy due to incorporation of annotation data, evidenced by the same analysis using a random annotation as negative control for a simulated trait (heritability = 0.5 or 0.1). SBayesRC_rand is SBayesRC when random numbers sampled from a uniform distribution between 0 and 1 are used as annotation data (a negative control for SBayesRC).

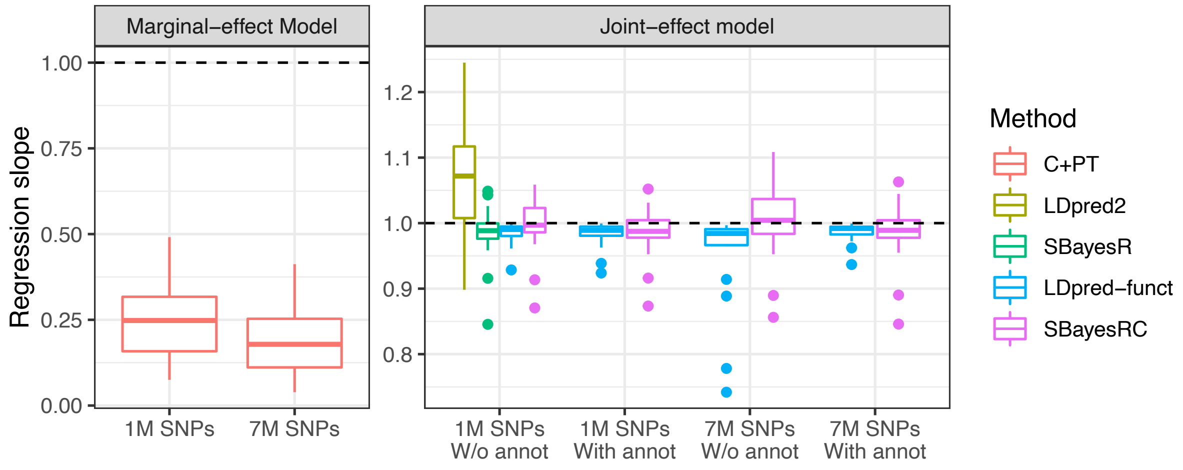


**Supplementary Figure 12** Mean regression slope from different methods in 28 independent traits across 10-fold cross validations in the UKB unrelated European sample. Note that 8 traits in LDPred-funct had a very large regression slope (> 5), hence were removed from the LDpred-funct column for better visibility of other methods. The values are shown in **Supplementary Table 3**.

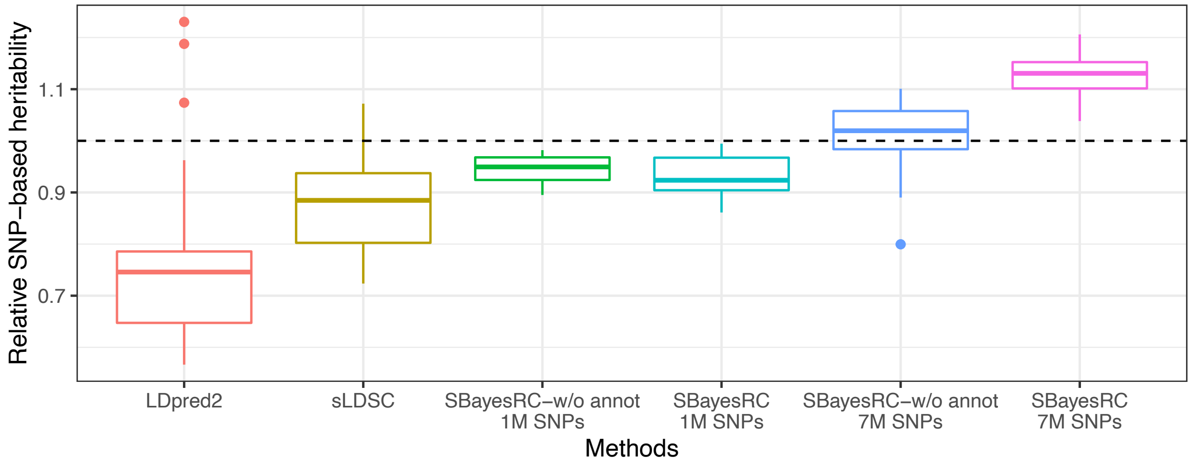


**Supplementary Figure 13** Relative SNP-based heritability estimate from different methods (baseline: estimate from SBayesR) for 28 independent traits in UKB unrelated European sample.

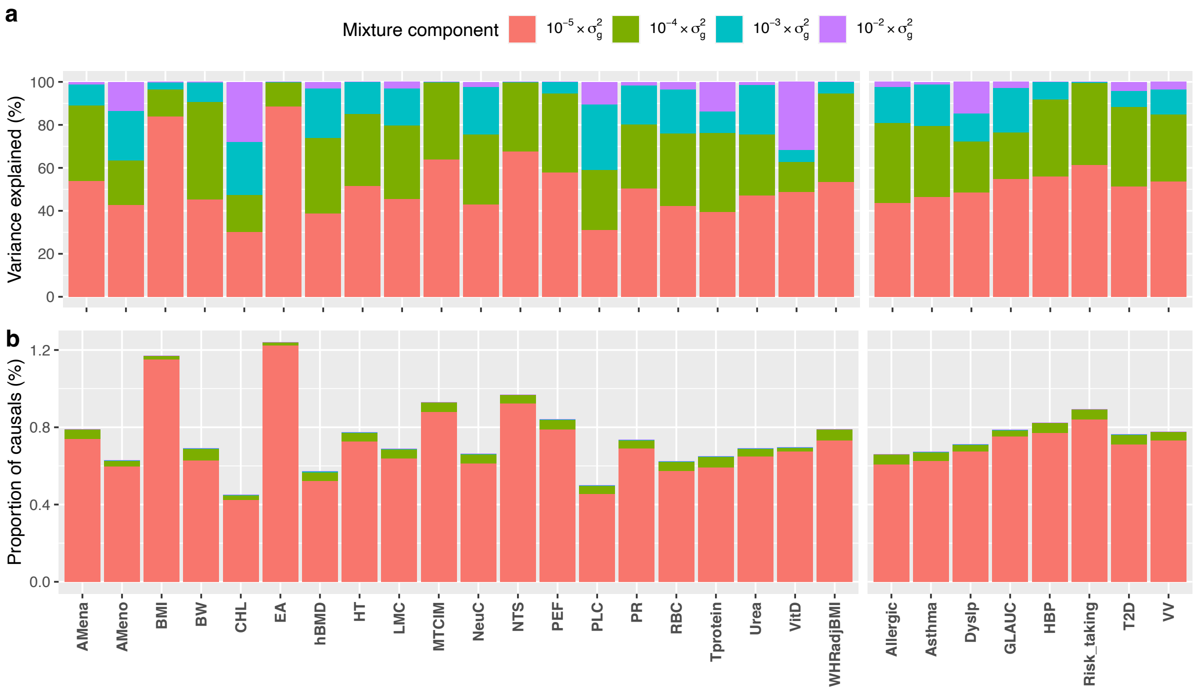


**Supplementary Figure 14** Genetic architecture estimates from SBayesRC using 7M common SNPs and annotation data for 28 independent traits in the UKB unrelated European sample. a) The proportion of genetic variance explained in the four non-zero mixture components. b) The proportion of causal variants allocated in the four non-zero mixture components.

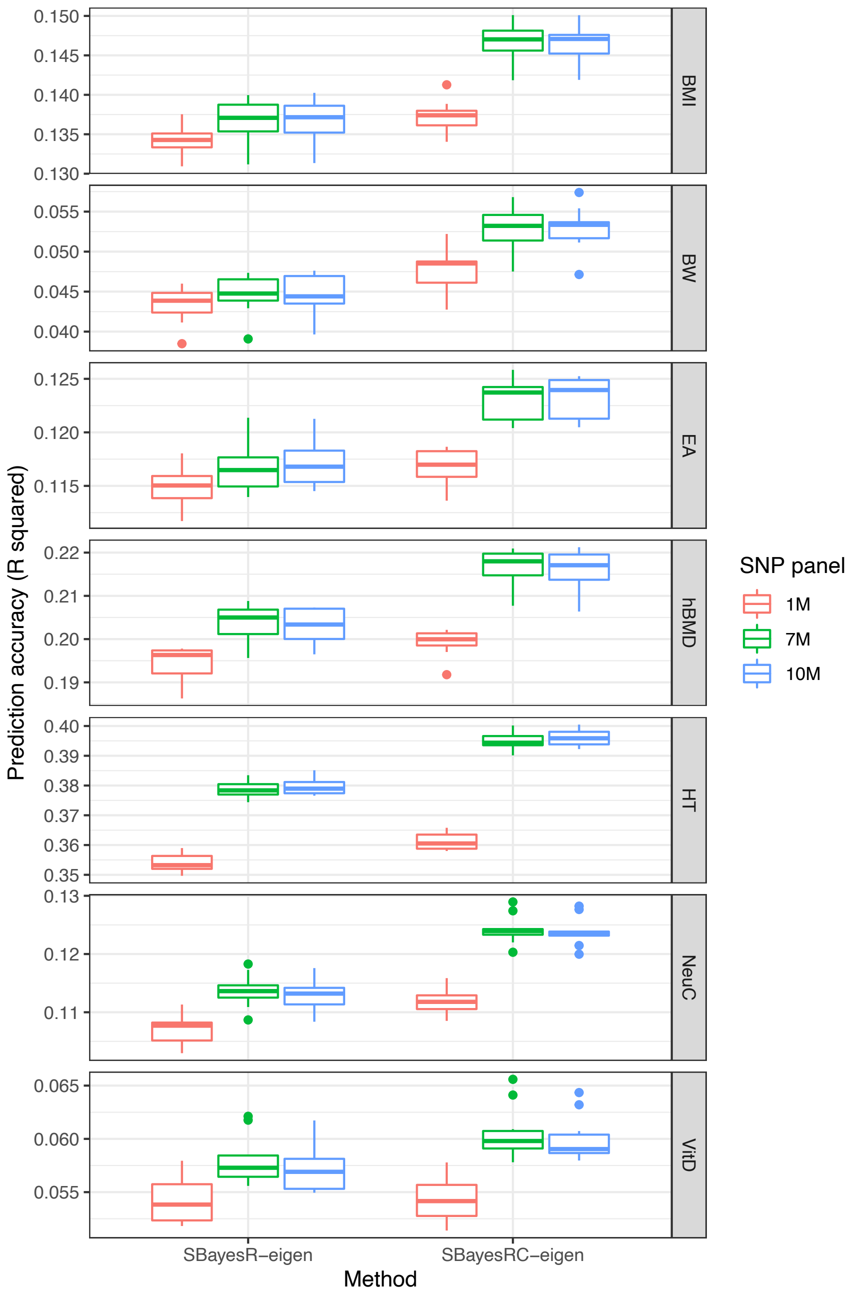


**Supplementary Figure 15** Prediction accuracy of SBayesRC without annotation or SBayesRC (incorporating annotation data) with 1M, 7M or 10M common SNPs for 7 UKB traits. Each box shows the results of 10-fold cross-validation in the unrelated European sample. Trait acronym BMI: body mass index; BW: birth weight; EA: educational attainment; hBMD: heel bone mineral density; HT: height; NeuC: neutrophile cell count; VitD: vitamin D level.

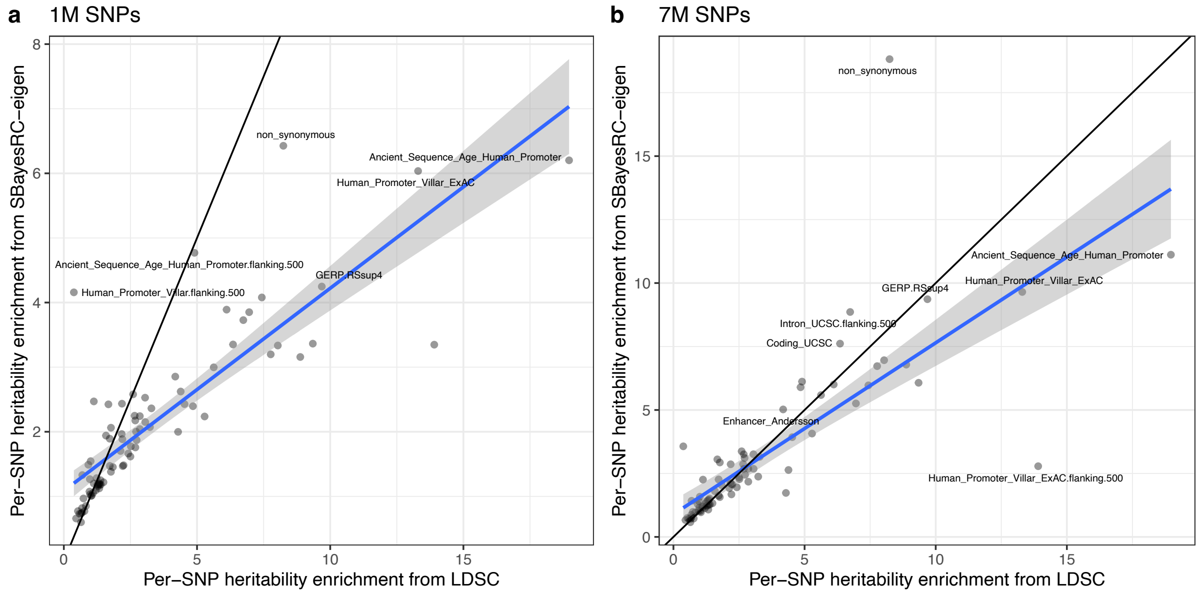


**Supplementary Figure 16** Comparison of per-SNP heritability enrichment in functional categories estimated by SBayesRC and S-LDSC using 1M (panel a) or 7M (panel b) SNPs, averaged over 28 independent traits from UKB.

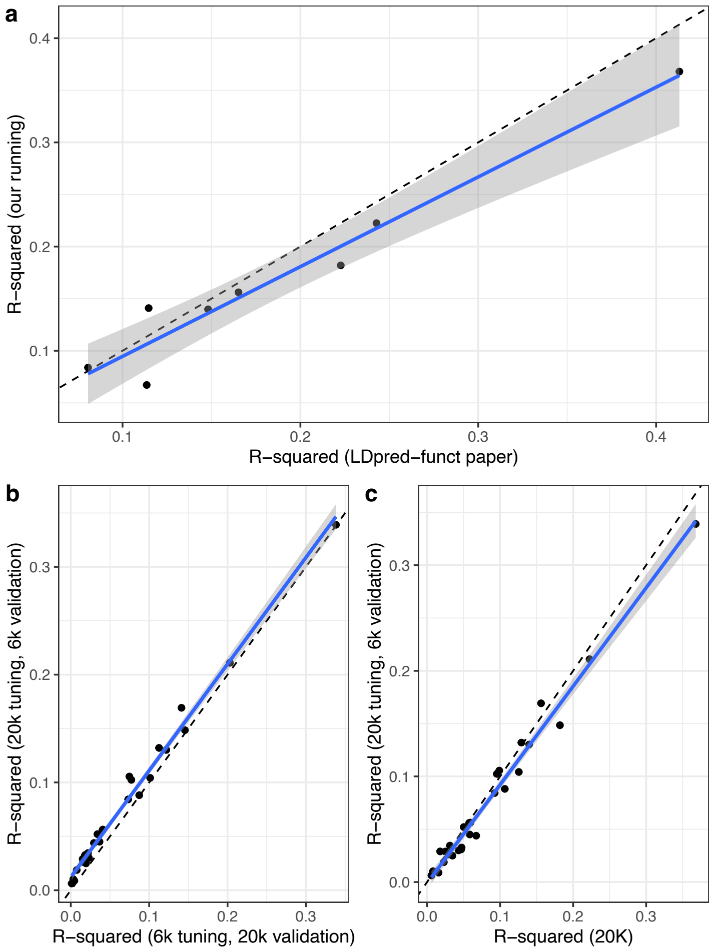


**Supplementary Figure 17** Comparison of prediction accuracy using LDpred-funct from this analysis to that reported in the LDpred-funct paper for the same trait. a) Comparison of prediction accuracy from our running (mean N=282,019) and that from the LDpred-funct paper (mean N=390,208). Although the prediction accuracies were highly correlated (r=0.975), we found a somewhat lower prediction accuracy in most traits, likely because of the smaller sample size used in this study. b) We found that a larger tuning sample size of 20K consistently gave better result than that of 6K in LDpred-funct. c) The default setting in LDpred-funct is to use the validation sample as tuning (x-axis), which gave slightly better prediction accuracy than using an independent validation sample but is subject to overfitting.

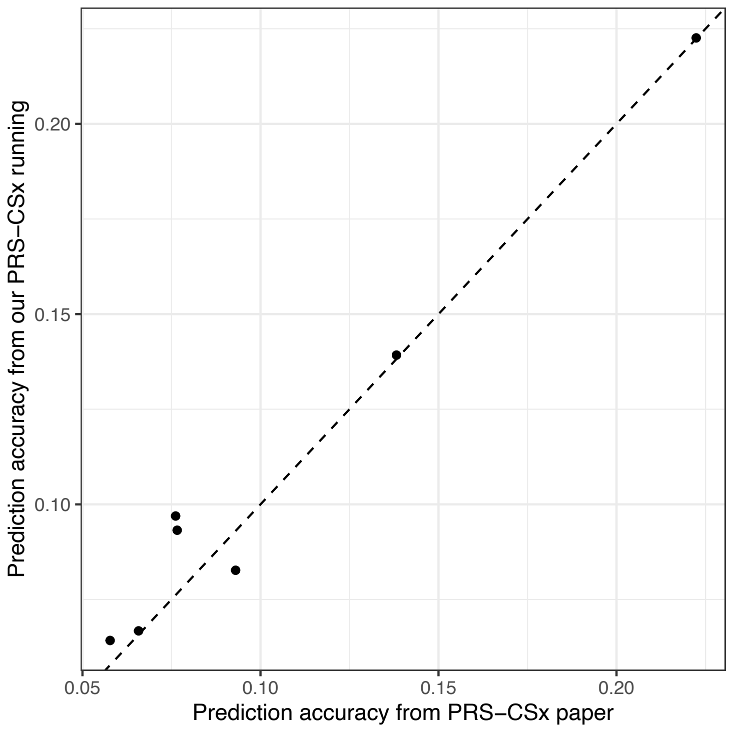


**Supplementary Figure 18** Comparison of prediction accuracy ($R^2$) using PRS-CSx from this analysis to that reported in the PRS-CSx paper for the same traits in EAS population training by summary data from UKB (EUR) and BBJ (EAS).

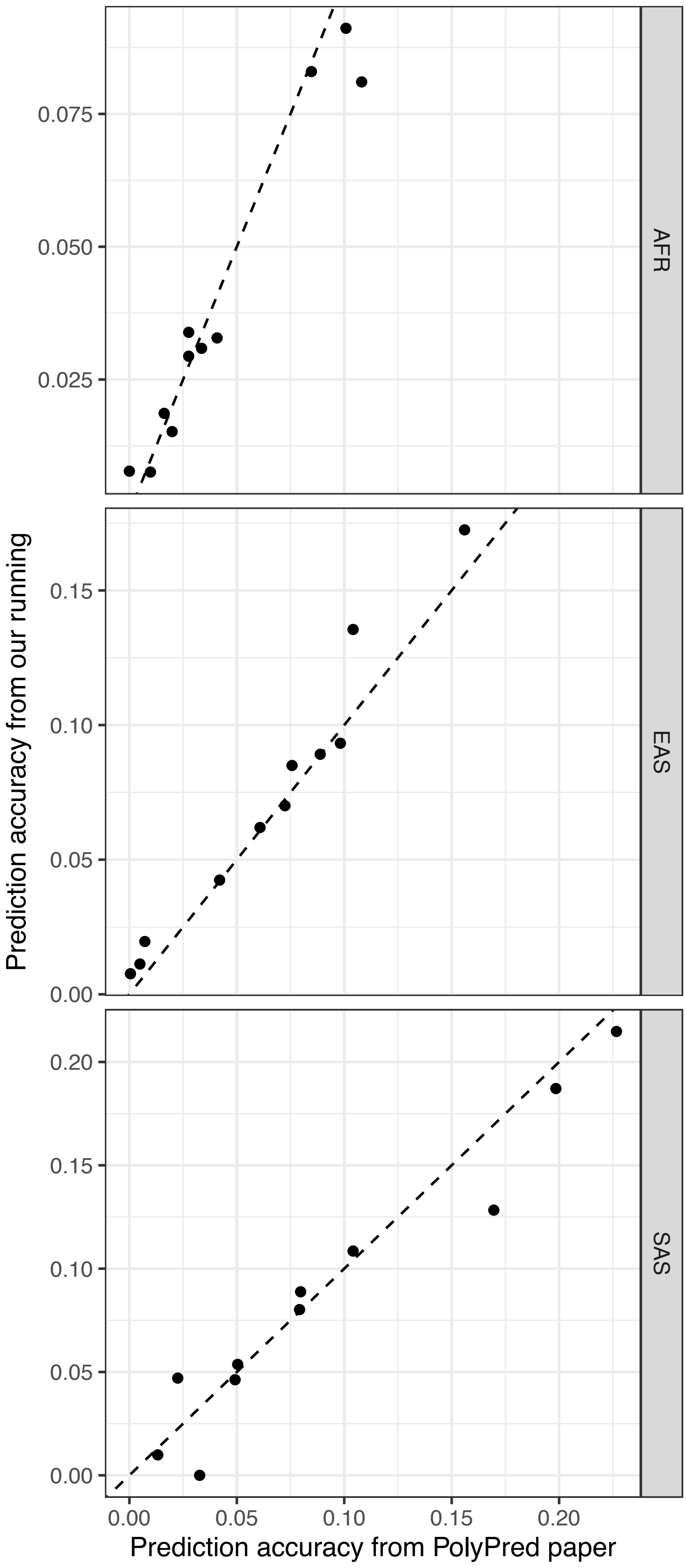


**Supplementary Figure 19** Comparison of prediction accuracy ($R^2$) using PolyPred-S from this analysis to that reported in the PolyPred paper for the same traits across ancestries. The prediction accuracies were highly correlated (r = 0.976). The differences may come from 1) Different ways to generate the summary statistics (linear regression/ logistic regression vs. linear mixed model); 2) Differences in training SNP panel (7 million vs. 18 million). We have also discussed with the authors to correct potential issues (https://github.com/omerwe/polyfun/issues/80).
